# GTP signaling links metabolism, DNA repair, and responses to genotoxic stress

**DOI:** 10.1101/2023.04.12.536297

**Authors:** Weihua Zhou, Zitong Zhao, Angelica Lin, John Yang, Jie Xu, Wilder-Romans Kari, Annabel Yang, Jing Li, Sumeet Solanki, Jennifer Speth, Natalie Walker, Andrew J. Scott, Ayesha U. Kothari, Yangyang Yao, Erik R. Peterson, Navyateja Korimerla, Christian K. Werner, Jessica Liang, Janna Jacobson, Sravya Palavalasa, Alexandra M. Obrien, Ameer L. Elaimy, Sean P. Ferris, Shuang G. Zhao, Jann N. Sarkaria, Balázs Győrffy, Shuqun Zhang, Wajd N Al-Holou, Yoshie Umemura, Meredith A. Morgan, Theodore S. Lawrence, Costas A. Lyssiotis, Marc Peters-Golden, Yatrik M. Shah, Daniel R. Wahl

**Affiliations:** Department of Radiation Oncology, University of Michigan, Ann Arbor, MI, USA; Department of Oncology, the Second Affiliated Hospital of Xi’an Jiaotong University, Xi’an, Shan Xi, PR China; Cell Signaling Technology, Inc., Danvers, MA, USA; Department of Molecular & Integrative Physiology, University of Michigan, Ann Arbor, MI, USA; Division of Pulmonary and Critical Care Medicine, Department of Internal Medicine, University of Michigan, Ann Arbor, MI, USA; Department of Oncology, the First Affiliated Hospital of Nanchang University, Nanchang, Jiangxi, PR China; Department of Pathology, Division of Neuropathology, University of Michigan, Ann Arbor, MI, USA; Department of Human Oncology, University of Wisconsin Madison, WI, USA; Mayo Clinic, Rochester, MN, USA; Department of Bioinformatics and 2(nd) Department of Pediatrics, Semmelweis University, Budapest, Hungary; TTK Cancer Biomarker Research Group, Institute of Enzymology, Budapest, Hungary; Department of Neurosurgery, University of Michigan, Ann Arbor, MI, United States; Department of Neurology, University of Michigan, Ann Arbor, MI, United States; Department of Molecular and Integrative Physiology and Department of Internal Medicine, Division of Gastroenterology and Hepatology, University of Michigan, Ann Arbor, MI, USA

**Keywords:** GTP signaling pathway, Rac1 pathway, Abl interactor 1(Abi-1), Phosphorylation and Dephosphorylation, NHEJ DNA Repair, Radioresistance, Radioprotection

## Abstract

How cell metabolism regulates DNA repair is incompletely understood. Here, we define a GTP-mediated signaling cascade that links metabolism to DNA repair and has significant therapeutic implications. GTP, but not other nucleotides, regulates the activity of Rac1, a G protein, that promotes the dephosphorylation of serine 323 on Abl-interactor 1 (Abi-1) by protein phosphatase 5 (PP5). Dephosphorylated Abi-1, a protein previously not known to activate DNA repair, promotes non-homologous end joining. In patients and mouse models of glioblastoma, Rac1 and dephosphorylated Abi-1 mediate DNA repair and resistance to standard of care genotoxic treatments. The GTP-Rac1-PP5-Abi-1 signaling axis is not limited to brain cancer, as GTP supplementation promotes DNA repair and Abi-1-S323 dephosphorylation in non-malignant cells and protects mouse tissues from genotoxic insult. This unexpected ability of GTP to regulate DNA repair independently of deoxynucleotide pools has important implications for normal physiology and cancer treatment.

## INTRODUCTION

Genotoxic therapy is a cornerstone of cancer treatment. In glioblastoma (GBM), the most common and aggressive primary brain cancer in adults, genotoxic radiation (RT) and temozolomide (TMZ) are standard of care treatments. The DNA damage response (DDR) controls the efficacy and toxicity of genotoxic therapy. The ability to rapidly repair RT- and TMZ-induced DNA damage causes treatment resistance and recurrence in GBM (1,2). Cancers with an impaired DDR are more sensitive to genotoxic agents such as RT (3), while patients with defective germline DDR have increased normal tissue toxicity from genotoxic therapy (4,5). Understanding the biology of the DDR has provided the opportunity to develop new drugs that could be used to augment the efficacy of genotoxic therapies (6). Indeed, inhibitors of the DDR are currently being investigated in combination with genotoxic agents for numerous cancers including GBM (7,8), though the risk for increased normal tissue toxicity from this approach could limit its efficacy.

Metabolism regulates cellular phenotypes, including the DDR. Our group and others have found links between metabolic processes, the DDR and resistance to genotoxic therapies such as RT and TMZ (9-13). Metabolites can regulate phenotypes through diverse mechanisms including regulating reactive oxygen species, driving macromolecule synthesis, activating signaling pathways and providing chemical energy. The molecular links between metabolites and the DDR have only been elucidated in a handful of cases. Metabolites that promote antioxidant synthesis can mitigate RT-induced toxicity by preventing oxidative DNA damage (14). Drugs such as gemcitabine and 5-FU deplete deoxynucleotides, cause cell cycle arrest, replication stress and increased RT efficacy. Both drugs are clinically used in combination with RT to treat a variety of cancers (15,16). More recently, groups have discovered that oncometabolite can impair homologous recombination by disrupting chromatin methylation at sites of DNA damage. This detailed understanding has led to clinical trials using PARP inhibitors, which are selectively efficacious in cancers lacking homologous recombination, in combination with genotoxic agents for these cancers(9,17,18). A mechanistic understanding of how metabolites regulate the DDR thus can have direct implications for patients.

In the present work, we define therapeutically relevant molecular links between nucleotide metabolism and the DDR in cancers and normal tissues. We find that GTP, but not other purines or pyrimidines, is the critical nucleotide that controls the DDR, which it does by regulating non-homologous end joining (NHEJ). Because only GTP regulated NHEJ, we reasoned that GTP-dependent signaling might be responsible for this regulation, rather than the ability for GTP to serve as a precursor for RNA or DNA, which it shares among other nucleotides. Using phosphoproteomics and a variety of molecular biology techniques and targeted assays, we identify a GTP-dependent dephosphorylation event on the protein Abl-interactor 1 (Abi-1) that regulates NHEJ. The dephosphorylation of Abi-1, a protein with no prior known role in DNA repair, depends on the activity of the G protein Rac1 and protein phosphatase 5. Modulating both Rac1 and Abi-1 activity affects the sensitivity of GBM to genotoxic therapies. These observations extend beyond GBM to normal tissues, as GTP supplementation promotes Abi-1 dephosphorylation in non-transformed cells and protects against normal tissue toxicity induced by RT and genotoxic chemotherapy. This surprising ability of GTP to regulate DNA repair independently of deoxynucleotide pools has important implications for normal physiology and the treatment of cancer.

## RESULTS

### GTP Promotes DSB Repair through NHEJ

We previously found that purines, but not pyrimidines, promote double-strand breaks (DSBs) repair after RT in GBM (11). The purine responsible for this regulation and its mechanistic details are unknown. To begin to understand these links, we first supplemented patient-derived neurosphere and immortalized GBM cell lines with the cell-permeable GTP precursor guanosine or the ATP precursor adenosine and assessed DSB repair after RT by measuring γ-H2AX (S139) foci formation and resolution by immunofluorescence (IF) as described previously (11). We found that guanosine, but not adenosine, supplementation decreased γ-H2AX foci at 4 h following RT in GBM neurosphere (Fig. 1A & S1A) and immortalized cell lines (Fig. 1B & S1B-D). Furthermore, silencing the rate-limiting enzymes of GTP (IMPDH1 and IMPDH2) but not ATP (ADSS1 and ADSS2) synthesis slowed DSB repair in both neurosphere and immortalized GBM cell models, which could be rescued by guanosine supplementation (Fig. 1C; Fig. S1E & F). Pharmacologic inhibition of purine synthesis by AG2037, which inhibits both GTP and ATP synthesis, slowed the repair of RT-induced DSBs. This activity was rescued by GTP (but not ATP) supplementation (Fig. 1D & S1G). Thus, we identified GTP as the key nucleotide responsible for DSB repair after RT in GBM models.

**Figure 1.**
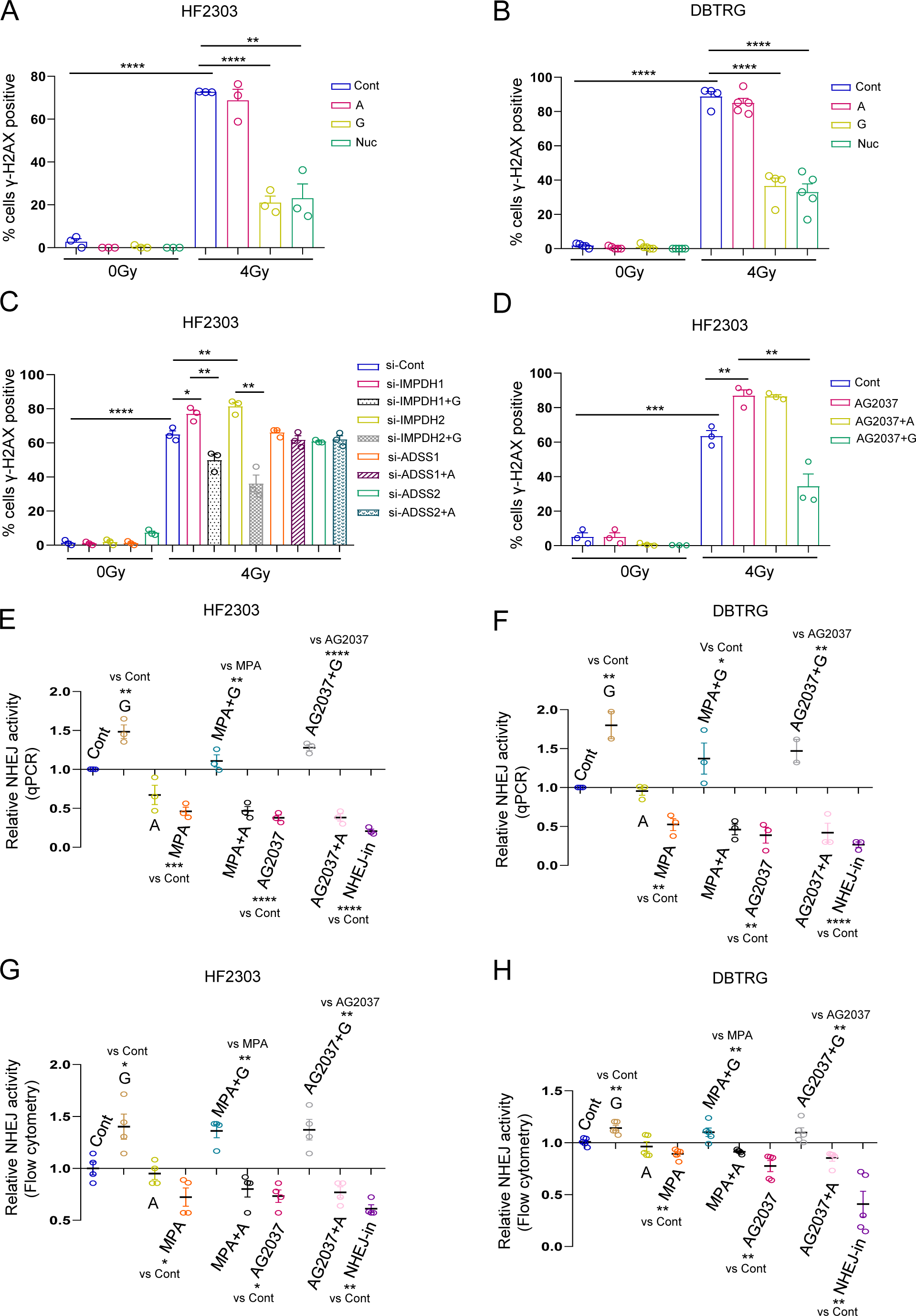
GTP promotes DSB repair through NHEJ. (A&B) Cells of the indicated line were treated with adenosine (A; 50 μm), guanosine (G; 50 μm) or pooled purine and pyrimidine nucleosides (Nuc; 8x) for 24 h and retreated with the same doses of individual purine or pooled nucleosides 2 h before RT and harvested for γ-H2AX (S139) quantification by IF 4 h post-RT. (C) Cells were transfected with a SMARTpool mixture of 4 siRNAs to target key enzymes of GTP or ATP synthesis and then treated with individual purine supplement and/or radiation as above, followed by γ-H2AX quantification by IF. (D) Cells were treated with AG2037 (150 nm) alone or combination with purine supplements (A or G). Cells were harvested 4 h post-RT for γ-H2AX quantification by IF. (E&F) Cells of the indicated line were treated with A (50 μm), G (50 μm), the GTP-only inhibitor mycophenolic acid (MPA, 10 μm), the dual GTP/ATP inhibitor AG2037 (150 nm) or their combinations and NHEJ was assessed using the qPCR-based pEYFP NHEJ system and normalized to untreated cells. The DNAPK inhibitor M3814 was used as a positive control (NHEJ-in). (G&H) Cells expressing the I-Scel-NHEJ reporter construct were treated with conditions identical to panels E&F and DNA damage was induced by infection with I-Scel adenovirus and NHEJ was assessed by quantifying the percentage of cells GFP positive 48 h later and normalized to untreated cells. Data are presented as mean ± SEM from three biologically independent experiments. Two-tailed unpaired student’s t test *p < 0.05, **p < 0.01, ***p < 0.001, ****p < 0.0001.

DSBs are primarily repaired through either non-homologous end joining (NHEJ) or homologous recombination (HR). GTP, but not ATP, supplementation enhanced NHEJ as assessed by both qPCR- and flow cytometry-based assays in multiple cell lines. Both GTP depletion by the IMPDH inhibitor mycophenolic acid (MPA) and combined GTP/ATP depletion by AG2037 decreased NHEJ repair as assessed by both assays. GTP, but not ATP, supplementation rescued the effects of both inhibitors (Fig. 1E-H). Modulating GTP levels did not affect HR activity (Fig. S1H) or the formation of RT-induced Rad51 foci, a marker of HR (Fig. S1I & J). Thus, we conclude that GTP but not ATP promotes DSB repair by promoting NHEJ.

### Dephosphorylation of Abi-1-S323 Is Crucial to GTP-induced NHEJ

GTP was unique among nucleotides in promoting NHEJ, therefore we reasoned that a property distinct from its ability to contribute to RNA or DNA synthesis was responsible for this link. Distinct from other nucleotides, GTP can activate unique signaling pathways through its ability to activate guanine nucleotide-binding proteins (G proteins) or through its conversion to cyclic GMP. To determine if GTP-specific signaling linked purine levels to NHEJ, we performed a phosphoproteomic assay in patient-derived GBM neurospheres to identify phosphorylation or dephosphorylation events that were induced by DNA damage (RT), augmented by GTP supplementation (G), and blocked by GTP depletion with mycophenolic acid (MPA, Fig. 2A). We identified approximately 17,000 phosphorylation sites, 2182 of which were candidate RT-induced phosphorylation events and 1913 candidate RT-induced dephosphorylation events. A single RT-induced phosphorylation site was potentiated by GTP supplementation but blocked by GTP depletion with MPA, while five RT-induced dephosphorylation sites were augmented by GTP supplementation but blocked by GTP depletion with MPA (Fig. 2A). Of these six candidate sites, we prioritized serine 323 on Abl interactor 1 (Abi-1, Fig. 2B), which has no known role in DNA repair but does canonically bind to the small G protein Rac1 (19). Notably, Rac1 can respond to physiologic changes in GTP levels (20). Consistent with this notion, our phosphoproteomic data suggested that the dephosphorylation of Abi-1 S323 that was blocked by GTP depletion with MPA treatment and was restored when GTP pools were refilled using guanosine supplementation (Fig. 2B). Thus, we hypothesized that dephosphorylation of S323 on Abi-1 might be a GTP-regulated signaling event that regulates DNA repair.

**Figure 2.**
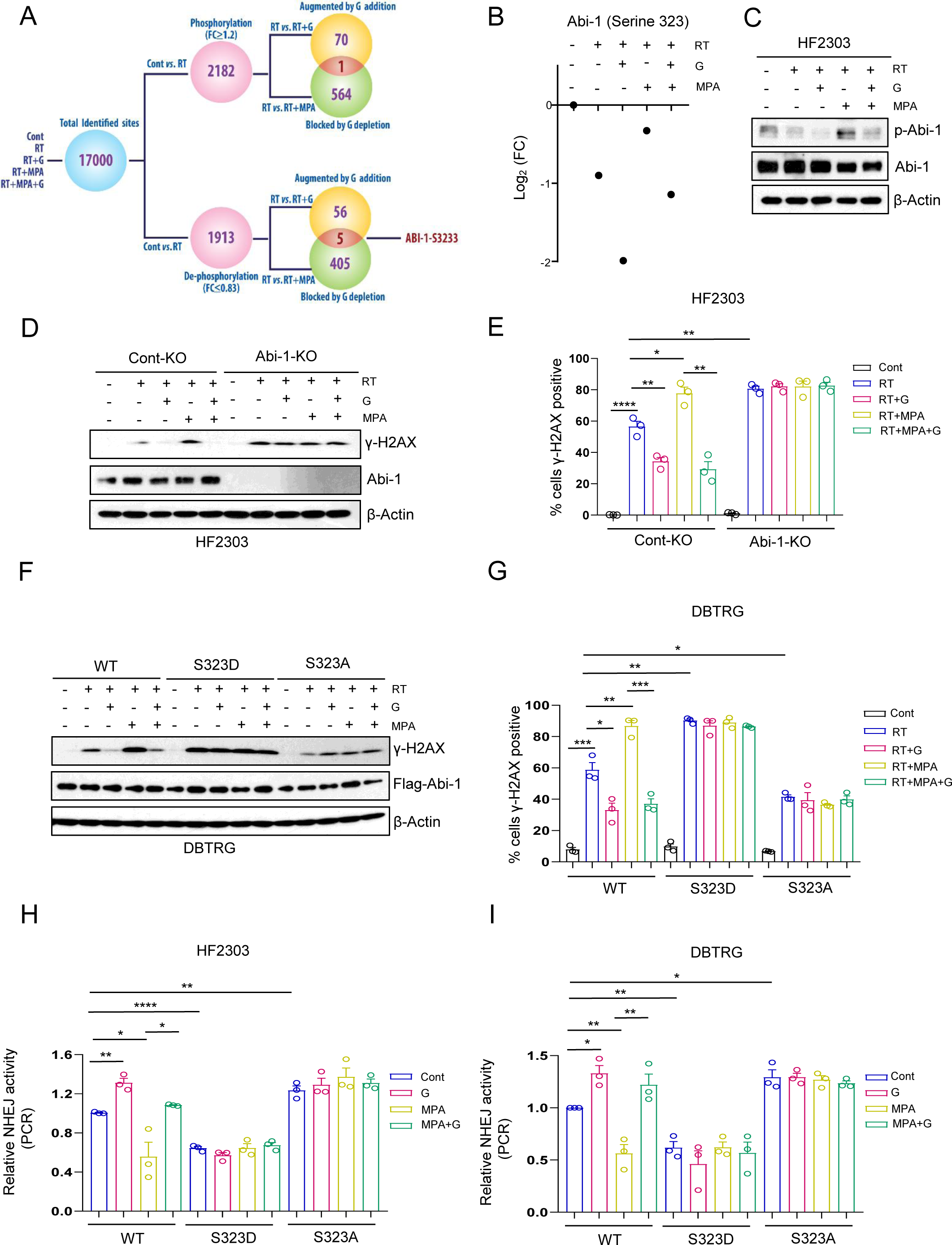
Dephosphorylation of Abi-1-S323 is crucial to GTP-induced NHEJ. (A) Schematic of phosphoproteomic assay, analysis pipeline and nomination of Abi-1-S323. (B) Levels of p-Abi-1-S323 as determined by phosphoproteomic assay. There is a DNA damage-induced dephosphorylation of Abi-1-S323 that is augmented by GTP supplementation but blocked by GTP deprivation with mycophenolic acid (MPA). The effects of MPA are abrogated when GTP supplementation is combined with MPA treatment. (C) Cells were treated with MPA (10 μm) and/or G (50 μm) for 24 h and retreated with G (50 μm) 2 h before RT, followed by cell harvesting 4 h after RT for immunoblot assay. (D&E) Control knockout (Cont-KO) or Abi-1 knockout (Abi-1-KO) cells were treated with MPA, G, and/or RT as before and harvested to assess γ-H2AX levels by immunoblot (D) or foci quantification (E). (F&G) Abi-1-KO cells were transfected with plasmids encoding wild type Abi-1, phospho-mimetic Abi-1 (S323D) or dephospho-mimetic Abi-1 (S323A) and treated as above, followed by immunoblot (F) or γ-H2AX foci assay (G). (H&I) Abi-1-KO cells were transfected with individual Abi-1 plasmid and treated with MPA and/or G overnight and retreated with G 2 h before transfection of linearized pEYFP products, followed by cell harvesting 24 h post transfection and qPCR assay with data normalized to untreated control cells. Data are presented as mean ± SEM from three biologically independent experiments for Figure E, G, H, I, and Figure C, D, F are representative figures from three biologically independent experiments. Two-tailed unpaired student’s t test *p < 0.05, **p < 0.01, ***p < 0.001, ****p < 0.0001.

To further interrogate this biology, we generated a phospho-Abi-1-S323 (p-Abi-1-S323) antibody and validated its specificity via western blot. The antibody showed a band at the expected molecular weight for Abi-1 in cells expressing wild type and phospho-mimetic Abi-1 (S323D), but not in cells expressing dephospho-mimetic Abi-1 (S323A) or with Abi-1 knocked out (Fig. S2A). We confirmed our phosphoproteomic data using this p-Abi-1-S323 antibody by showing that p-Abi-1-S323 was decreased after RT and could be decreased further after guanosine supplementation but blocked by GTP depletion (Fig. 2C & S2B).

To interrogate the role of Abi-1 and its dephosphorylation in the DDR, we generated Abi-1-knockout (KO) GBM cell lines and neurospheres using CRISPR-CAS9 (Fig. 2D). DSB repair was evaluated by immunoblot to detect γ-H2AX protein level and IF to detect γ-H2AX foci after DNA damage with RT. Consistent with our previous study (11) and initial findings (Fig. 1), repair of RT-induced DSBs was promoted by GTP supplementation but blocked by GTP depletion in control-KO GBM cells. Cells with Abi-1-KO had increased γ-H2AX following RT suggestive of slow DSB repair. Furthermore, the amount of γ-H2AX in these cells following RT was no longer influenced by GTP supplementation or depletion (Fig. 2D & E; S2C & D). These data suggest that Abi-1 plays a role in GTP-mediated DNA repair. To specifically interrogate the dephosphorylation of Abi-1-S323, we re-expressed WT, S323D/phospho-mimetic, or S323A/dephospho-mimetic Abi-1 into Abi-1-KO cells and evaluated γ-H2AX levels following RT by immunoblot and foci formation. Re-expression of phospho-mimetic Abi-1 (S323D) slowed but dephospho-mimetic Abi-1 (S323A) enhanced DSB repair. GTP supplementation or depletion only influenced DSB repair in cells expressing Abi-1-WT, but not in those expressing either point mutant (Fig. 2F & G; S2E & F).

To examine which DSB pathway was regulated by Abi-1 dephosphorylation, we employed the linearized plasmid-based end-joining assay to interrogate NHEJ and the RAD51 foci assay to interrogate HR. Cells expressing Abi-1-S323A had enhanced NHEJ while those expressing Abi-S323D had impaired NHEJ. Further, NHEJ was no longer influenced by GTP supplementation or depletion in cells expressing these point mutants (Fig. 2H & I). Mutating Abi-1-S323 did not affect HR activity (Fig. S2G & H). Together, these findings indicate that the GTP-induced dephosphorylation of Abi-1-S323 is critical for DSB repair through NHEJ. Furthermore, because GTP modulation no longer affected DSB repair or NHEJ in cells expressing mutated Abi-1 S323, our data suggest that the main role of GTP in promoting NHEJ and DSB repair is through promoting Abi-1 dephosphorylation rather than as acting as a physical component of newly synthesized nucleic acids.

### Rac1 Controls the GTP-dependent Dephosphorylation of Abi-1-S323

We next sought to find out how changes in GTP levels cause the dephosphorylation of Abi-1-S323 to promote DNA repair. Abi-1 canonically binds to the G protein Rac1 (19) and promotes actin cytoskeleton remodeling and cell mobility (21,22). To determine if Rac1 promotes Abi-1-S323 dephosphorylation and subsequent DNA repair, we introduced three Rac1 plasmids (Rac1-WT, dominant negative mutant Rac1-T17N and active mutant Rac1-Q61L) and confirmed their activity using a Rac1 activity assay (Fig. S3A). While GTP supplementation with its precursor guanosine increased Rac1 activity after radiation, GTP depletion with MPA blocked Rac1 activity in cells expressing the Rac1-WT plasmid. Cells expressing constitutively active Rac1-Q61L mutant maintained high Rac1 activity, whereas those expressing dominant negative -T17N mutant showed minimal Rac1 activity after RT. Rac1 activity was not modified by GTP supplementation or depletion in cells expressing Rac1-Q61L or Rac1-T17N (Fig. 3A & S3B). Furthermore, cells expressing dominant negative Rac1-T17N had higher levels of γ-H2AX following RT compared to cells expressing Rac1-WT, while those expressing Rac1-Q61L had lower levels (Fig. 3B & S3C). Supplementing or depleting GTP levels no longer influenced γ-H2AX levels following RT in cells expressing constitutively active or dominant negative Rac1. These data suggest that GTP-dependent Rac1 activity is important for RT-induced DSB repair.

**Figure 3.**
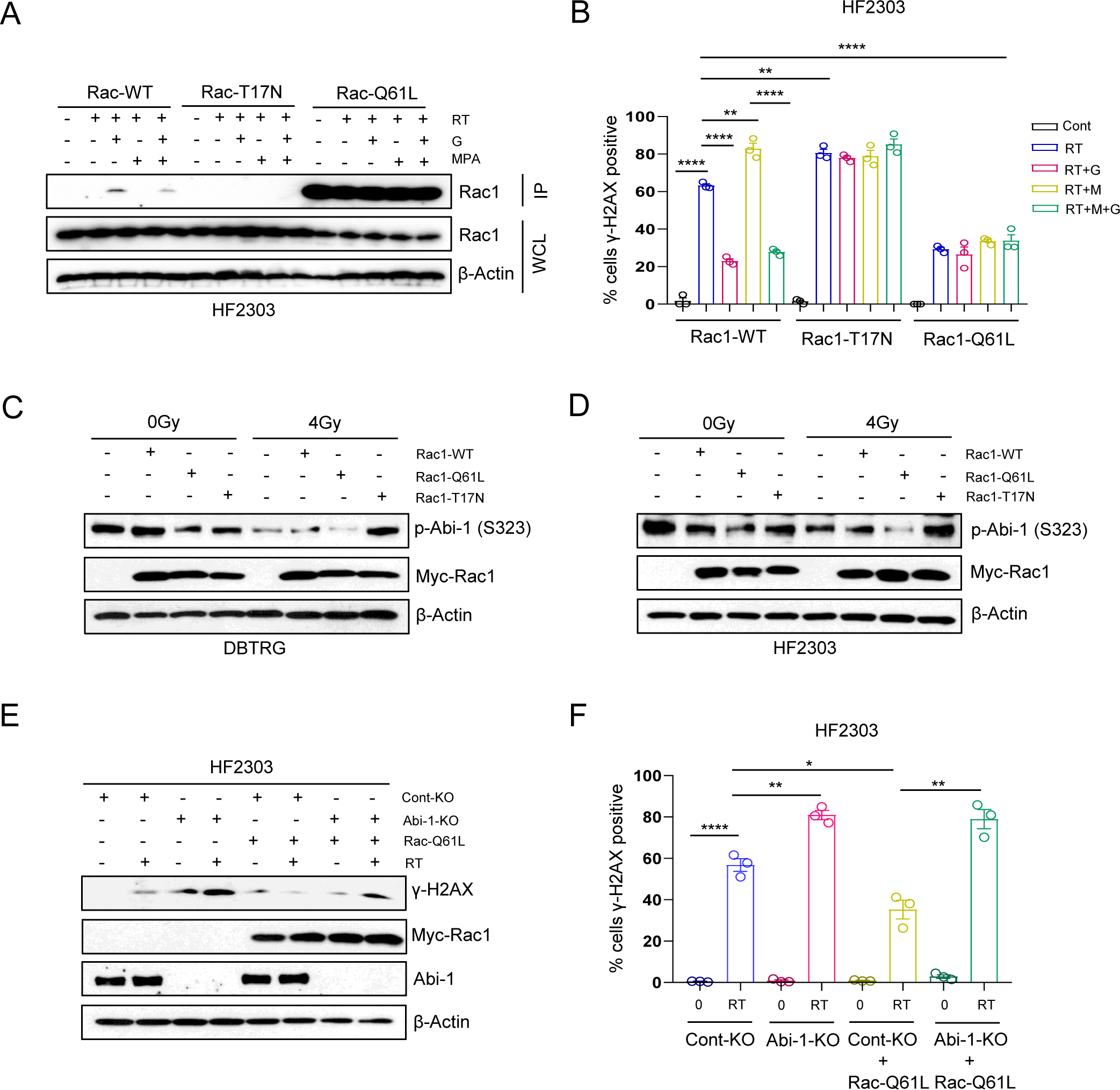
Rac1 controls the GTP-dependent dephosphorylation of Abi-1-S323. (A&B) Cells were transfected with plasmids encoding Rac1-WT, -T17N (dominant negative), or - Q61L (constitutively active), and treated with MPA, G, or RT as in Figure 2. Cells were harvested 4 h post RT to assess Rac1 activity (A) or γ-H2AX foci by immunofluorescence (B). (C&D) Cells were transfected with indicated Rac1 plasmids (WT, constitutively active Q61L or dominant negative T17N) and irradiated with 4 Gy, followed by cell harvesting 4 h post-RT to assess p-Abi1-S323 by immunoblot. (E&F) Con-KO or Abi-KO cells were transfected with Rac1-Q61L plasmid and treated with radiation and harvested for immunoblot (E) or γ-H2AX foci IF staining (F). Data are presented as mean ± SEM from three biologically independent experiments for Figure B & F and Figure A, C, D, E are representative figures from three biologically independent experiments. Two-tailed unpaired student’s t test *p < 0.05, **p < 0.01, ****p < 0.0001.

To investigate whether the dephosphorylation of Abi-1-S323 occurs downstream of Rac1 activity, we conducted epistasis experiments. Irradiating cells expressing Rac1-WT caused the dephosphorylation of Abi-1-S323. This desphosphorylation was augmented in cells expressing constitutively active Rac1-Q61L but was blocked in cells expressing dominant negative Rac1-T17N (Fig. 3C & D). Furthermore, expression of constitutively active Rac1-Q61L no longer promoted DSB repair following RT in cells lacking Abi-1, as assessed by both γ-H2AX protein level (Fig. 3E & S3D) and foci levels (Fig. 3F & S3E). Thus, our data confirmed that GTP-activated Rac1 promotes dephosphorylation of Abi-1-S323, which in turn promotes the repair of RT-induced DSBs.

### Protein Phosphatase 5 Mediates the GTP/Rac1-Depdendent Dephosphorylation of Abi-1 (S323) and Downstream DSB repair

We next sought to identify the phosphatase responsible for mediating the Rac1-dependent dephosphorylation of Abi-1-S323. We treated GBM cells (DBTRG) or neutrospheres (HF2303) with okadaic acid (OA), which inhibits protein phosphatase (PP)1, 2A, 4, and 5, and fostriecin (Fos), which inhibits PP2A and 4 (23). We found that the RT-induced dephosphorylation of Abi-1-S323 was blocked by okadaic acid but not fostriecin (Fig. 4A & S4A). This finding suggested that PP1 or PP5 might be responsible for Rac1-dependent Abi-1-S323 dephosphorylation. To further elucidate the responsible phosphatase, we individually silenced PP1, PP2A, PP4, and PP5, and found that only silencing PP5 blocked the RT-induced decrease of p-Abi-1-S323 (Fig. 4B & S4B). These data indicate that PP5 is responsible for Abi-1-S323 dephosphorylation after DNA damage and are consistent with previous reports that Rac1 can bind to and activate phosphatase 5 (PP5) (24,25).

**Figure 4.**
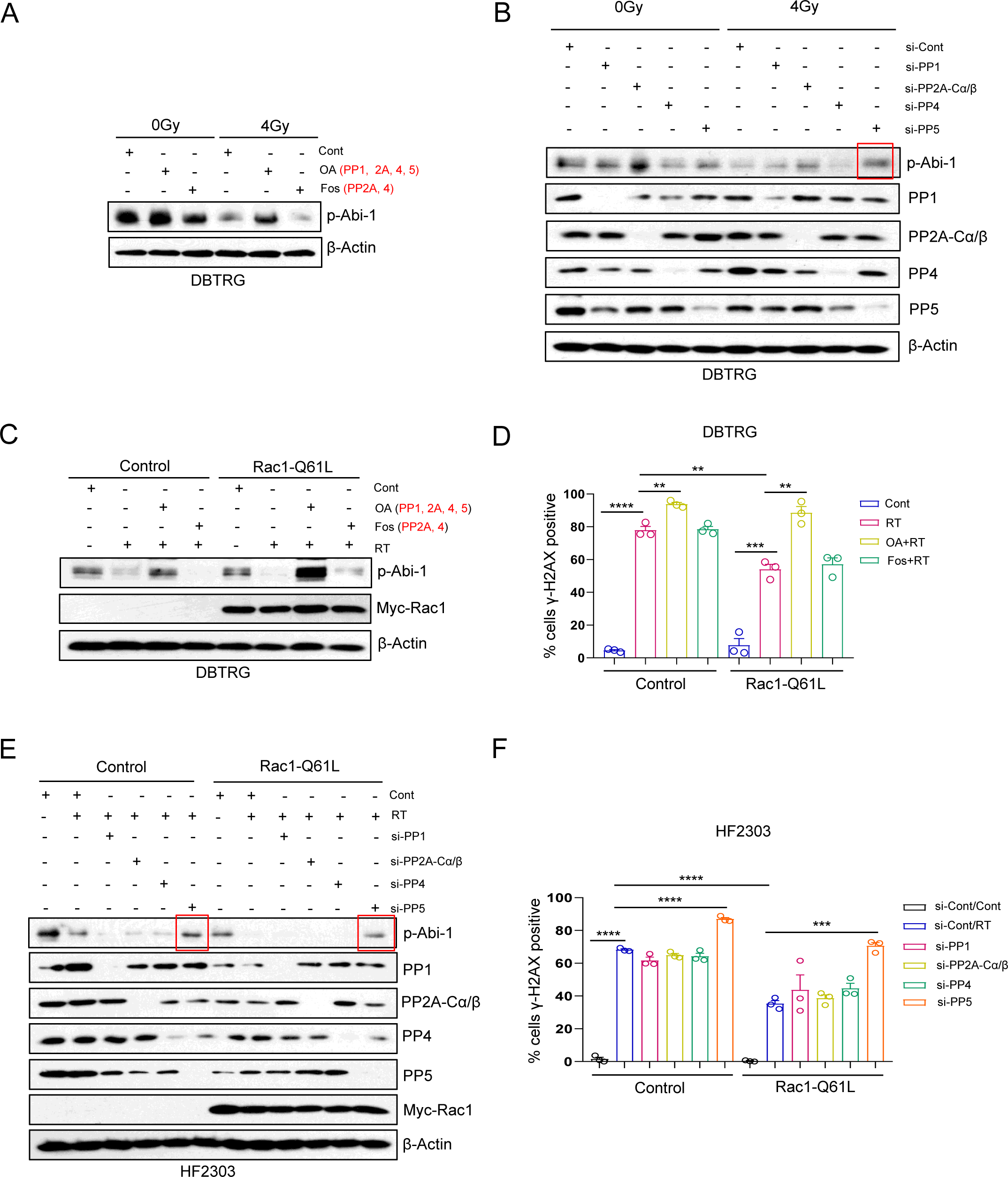
Protein phosphatase 5 mediates the GTP/Rac1-depdendent dephosphorylation of Abi-1 (S323) and downstream DSB repair. (A&B) Cells were treated with the phosphatase inhibitors okadaic acid (OA; 15 nm) or fostricein (Fos; 100 nm) 1 h before RT (A), or first transfected with pan PP1 siRNA, mixture of PP2A catalytic (PP2A-C) subunit α/β, PP4 and PP5 for 48 h and irradiated after transfection (B), followed by cell harvesting 4 h post-RT for immunoblot. (C-F) Cells were transfected with constitutively active Rac1-Q61L and then treated with okadaic acid (15 nm) or fostricein (100 nm) 1 h before RT (C&D), or transfected with individual phosphatase siRNA pool, along with overexpression of Rac1-Q61L or control (E&F), followed by immunoblot (C&E) or γ-H2AX foci IF staining (D&F). Data are presented as mean ± SEM from three biologically independent experiments for Figure D, F and Figure A, B, C, E are representative figures from three biologically independent experiments. Two-tailed unpaired student’s t test **p < 0.01, ***p < 0.001, ****p < 0.0001.

We next determined whether the PP5-induced dephosphorylation of Abi-1-S323 is GTP-Rac1-dependent and responsible for DSB repair. Expression of constitutively active Rac1-Q61L augmented the dephosphorylation of Abi-1-S323 and enhanced DSB repair following RT. These effects were blocked by okadaic acid, but not fostriencin (Fig. 4C & D; Fig. S4C & D). Likewise, we confirmed that only silencing PP5 could blocked the augmented DSB repair found in cells expressing constitutively active Rac1-Q61L (Fig. 4E & F; S4E & F). Our data indicate the presence of a signaling axis in which high GTP levels promote Rac1 activity, which causes the PP5-mediated dephosphorylation of Abi-1-S323, which activates DSB repair through NHEJ.

### Rac1 Activity Influences GBM Treatment Responses

Increased Rac1 activity correlates with therapeutic resistance and shorter survival in many cancers, suggesting that it may be a promising target for cancer therapy (26). To investigate these associations in GBM, we confirmed that high transcript expression of Rac1 and its downstream targets (Table S1) is associated with inferior survival of GBM patients (Fig. S5A & B). We interrogated the DepMap and found that Rac1 is co-essential with Abi-1 across over 1000 cancer cell lines (Fig. S5C), suggesting that some of the oncogenic properties of Rac1 may be due to its ability to regulate Abi-1. To further demonstrate the role of Rac1 and p-Abi-1-S323 in GBM treatment resistance, we turned to the Mayo Clinic Brain Tumor Patient Derived Xenograft (PDX) National Resource, which has profiled the treatment responses of dozens GBM PDX models (27,28). Using a tissue microarray constructed from the GBM PDXs in this database, we measured phosphorylation levels of Abi-1-S323 and two other Rac1 downstream proteins, PAK 1 and PAK2, both of which are phosphorylated when activated by Rac1 (29) (30). We confirmed that our new p-Abi-1-S323 antibody was appropriate for immunohistochemistry (IHC) by testing it in flank and intracranial Abi-1 knockout GBM tumors and confirming absent signal in both models (Fig. S5D). In the PDX GBM tissue samples, we found that increased phosphorylation of PAK1 and PAK2 was correlated with resistance to both temozolomide (TMZ) and combined TMZ and RT treatment. By contrast, a lack of phosphorylated Abi-1-S323 was associated with TMZ and TMZ/RT treatment resistance (Fig. 5A-F).

**Figure 5.**
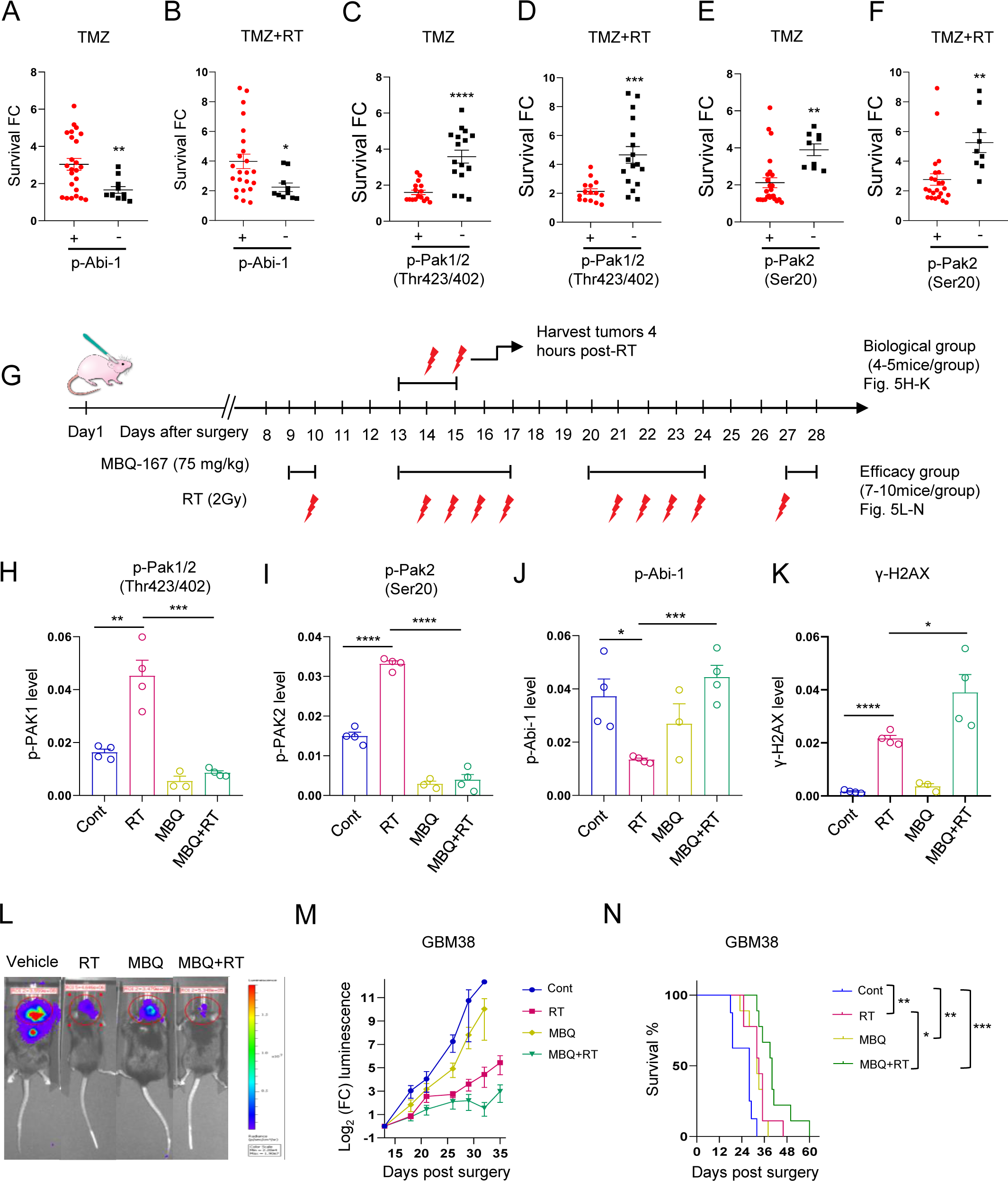
Rac1 activity influences GBM treatment responses. (A-F) IHC staining was performed to detect expression of p-Abi-1 and two downstream proteins of active Rac1 (PAK1 and PAK2) in GBM PDX tissue arrays. Survival FC indicates the fold increase in median survival with the indicated treatment compared to placebo control. Each dot indicates a different murine PDX model. (G) A schematic timeline of GBM38 orthotopic mouse models (additional details in Methods). (H-K) A subset of mice (3-4 mice/group) were treated with three doses of the Rac1 inhibitor MBQ-167 and two fractions (2 Gy/fraction) of radiation and tumors were harvested 4 h after receiving the second RT dose for IHC staining with indicated antibodies. (L-N) Another subset mice (7-10 mice/group) were treated with 12 doses of MBQ-167 and 10 fractions (2 Gy/fraction) of radiation (as shown in Figure 5G), were used for efficacy evaluation. Mice were treated with 150 mg/kg D-luciferin and imaged 10 min post-injection (L). Total flux of equal-area ROIs at each time point were normalized to flux at the first day of treatment for evaluating tumor progression (M). Mice were monitored daily and euthanized when they developed neurologic symptoms and Kaplan–Meier survival curve was plotted (N). Data are presented as mean ± SEM from 3-4 independent mice for Figure H-K and 7-10 mice for figure L-N. Two-tailed unpaired student’s t test **p < 0.01, ***p < 0.001, ****p < 0.0001 for Fig. 5A-F and H-K; Log-rank (Mantel-Cox) test *p < 0.05, **p < 0.01, ***p < 0.001 for Fig. 5N.

We next sought to intervene on this pathway to slow GBM DNA repair and overcome treatment resistance. We generated RT-resistant orthotopic GBM PDXs (GBM38) and treated them in four groups: RT alone; Rac1 inhibitor MBQ-167 (MBQ) alone; combined RT and MBQ-167; and vehicle treatment. To assess for target engagement, signaling and DNA repair, mice were treated with an abbreviated regimen of two daily fractions of RT (2 Gy/fraction) and three daily treatments of MBQ-167 (75 mg/kg/day) with tumors harvested 4 hours following their final RT treatment (Fig. 5G). RT increased the phosphorylation of PAK1 and PAK2 and decreased the phosphorylation of Abi-1-S323 (Fig. 5H-J & S5E). Importantly, inhibition of Rac1 by MBQ-167 stopped the RT-induced phosphorylation of PAK1 and PAK2 and dephosphorylation of p-Abi-1-S323. MBQ-167 treatment also slowed the repair of RT-induced DSBs as evidenced by increased γ-H2AX staining (Fig. 5K & S5E), which suggests that increased Rac1 activity post RT can promote dephosphorylation of Abi-1-S3323 and DSB repair.

To assess for slowing of GBM tumor growth, we treated tumor-bearing mice with 10 fractions of RT and/or 12 doses of MBQ-167 (Fig. 5G). RT alone slowed tumor growth and extended mouse survival, and its activity was augmented when RT was combined with the Rac1 inhibitor (Fig. 5L-N). Similar results were observed in a second orthotopic GBM model (HF2303, Fig. S5F-I). Thus, active Rac1 promotes DNA repair and RT resistance through dephosphorylation of Abi-1-S323 and inhibiting Rac1 can block this signaling and increase treatment efficacy.

### Abi-1 Mediates Genotoxic Treatment Efficacy in GBM

Having found that Abi-1 regulates DNA repair and that Rac1 activity influences GBM treatment resistance, we set out to determine if Abi-1 and its dephosphorylation regulated the response of GBMs to genotoxic treatments. Analysis of glioma cell lines in the DepMap indicated that Abi-1 expression is correlated with resistance to bleomycin, an anticancer drug that mediates cell death by inducing DSBs (31) (Fig. S6A). Furthermore, high Abi-1 expression correlated with resistance to TMZ (p = 0.00298) and TMZ+RT (p = 0.00006) in GBM PDXs (28) (Fig. S6B & C), which suggests that Abi-1 might directly protect GBMs against genotoxic treatments.

Using our Abi-1 knockout cell lines and neurospheres, we then set out to understand how Abi-1 regulated GBM responses to a variety of genotoxic therapies. Abi-1 knockout enhanced RT-sensitivity at numerous doses in both HF2303 neurospheres (Fig. 6A) and immortalized DBTRG cells (Fig. S6D). Furthermore, Abi-1 knockout enhanced the effects of bleomycin (Fig. 6B & S6E), TMZ (Fig. 6C & S6F), and TMZ+RT (4Gy) (Fig. 6D & S6G) by decreasing their IC50 in both cell models. To ensure that Abi-1 knockout selectively sensitizes GBMs to genotoxic treatments rather than more general cell death, we repeated these experiments with non-genotoxic chemotherapies. Abi-1 knockout did not affect cell sensitivity to paclitaxel or vincristine (Fig. S6H-K), which exhibit anti-cancer ability by targeting microtubes (32,33). To ensure that it was the dephosphorylation of Abi-1 that mediated resistance to genotoxic therapy, we re-expressed Abi-1-WT, -S323D and -S323A back into Abi-1-KO cells. Compared to Abi-1-WT, re-expression of the dephospho-mimetic Abi-1-S323A restored resistance to genotoxic therapies, while the phospho-mimetic mutant Abi-1-S323D enhanced their sensitivity (Fig. 6E-H; S6L-O), which suggests that dephosphorylation of Abi-1-S323 helps mediate GBM resistance to genotoxic therapies.

**Figure 6.**
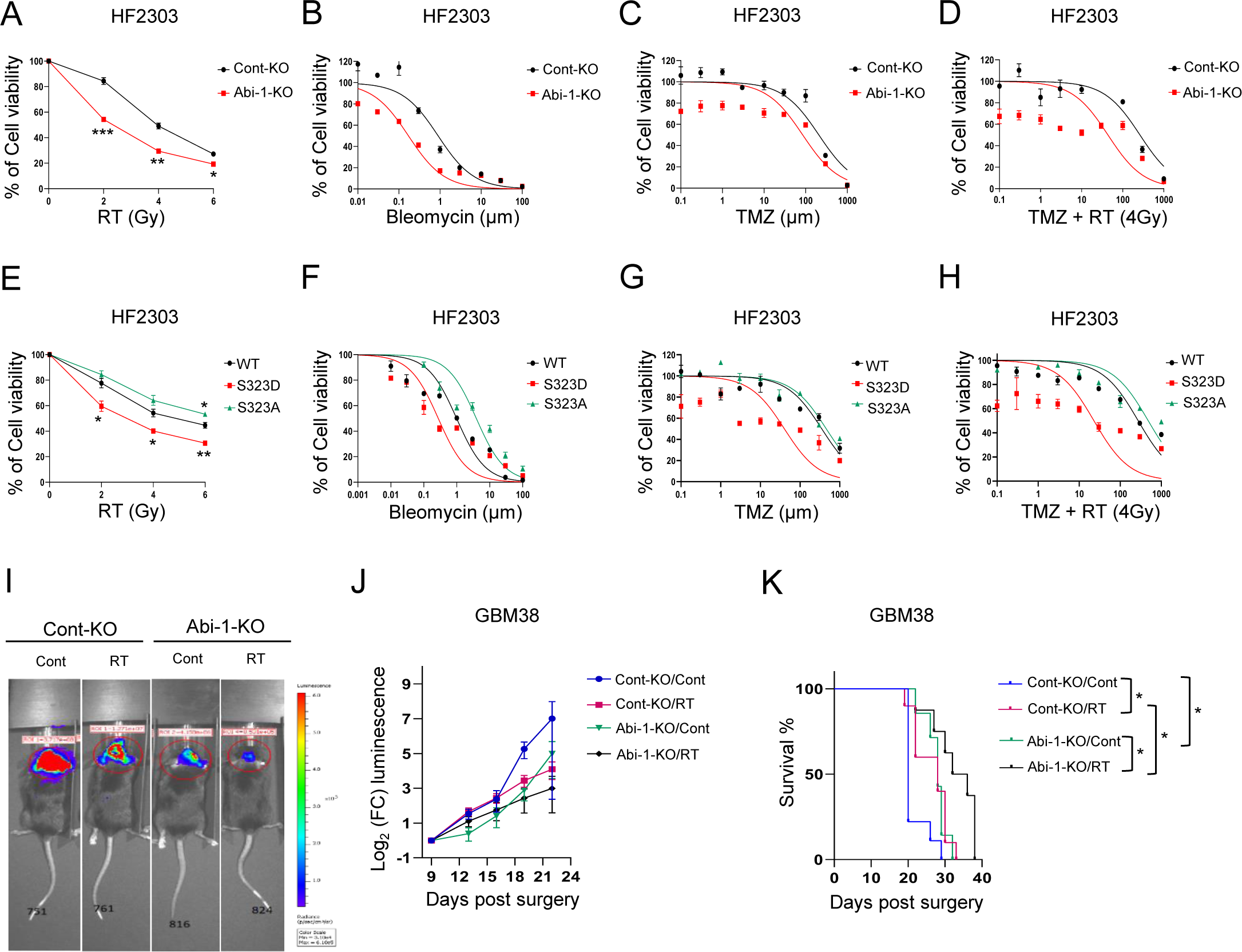
Abi-1 mediates genotoxic treatment efficacy in GBM. (A-D) Con-KO or Abi-1-KO cells were treated with different doses of radiation (A) or escalating concentrations of bleomycin (B; Con-KO vs Abi-KO IC50:0.77 ± 0.07 vs 0.30 ± 0.09, p = 0.02), TMZ alone (C; 265.33 ± 34.77 vs 134.60 ± 25.71, p = 0.04) or TMZ combined with RT (4 Gy) (D; 229.23 ± 41.65 vs 75.68 ± 21.61, p = 0.03) and cell viability was evaluated with long-term cell viability assay at day 7 post treatment. All drug concentrations are in μM. (E-H) Abi-KO cells were transiently transfected with Abi-1-WT, -S323D or -S323A, followed by irradiation (E), bleomycin (F; IC50 of WT vs S323D:1.13 ± 0.11 vs 0.28 ± 0.004, p = 0.03; WT vs S323A:1.13 ± 0.11 vs 4.32 ± 0.33, p = 0.02), TMZ alone (G; IC50 of WT vs S323D: 324.53 ± 16.43 vs 90.16 ± 38.91, p = 0.005; WT vs S323A: 324.53 ± 16.43 vs 425.73 ± 23.61, p = 0.02) or TMZ combined with RT (4 Gy) (H; IC50 of WT vs S323D:215.13 ± 46.12 vs 47.19 ± 24.61, p=0.03; WT vs S323A: 215.13 ± 46.12 vs 392.43 ± 44.64, p = 0.05) treatment as discussed above, all drug concentrations in μM. (I-K) Luciferase-positive, cont or Abi-1 knockout, and RT-resistant GBM38 patient-derived xenograft cells were orthotopically implanted and tumor-bearing mice were randomized (7-10 animals per group). RT (2 Gy/fraction and 6 fractions for total) was given. Tumors were imaged after D-luciferin injection and mouse survival was monitored (mean ± SEM). Figure A to H are representative figures from two to three biologically independent experiments (mean ± SEM) and I-K are presented as mean ± SEM from 7-10 independent mice. Two-tailed unpaired student’s t test *p < 0.05, **p < 0.01, ***p < 0.001, for Fig. 6A-H; Log-rank (Mantel-Cox) test *p < 0.05, for Fig. 5K.

To investigate this biology *in vivo*, we generated orthotopic GBM tumors using control or Abi-1 knockout GBM38 cells and gave the mice 6 fractions of radiation treatment once tumors formed (Fig. S6P). In control tumors, RT slowed tumor growth (Fig. 6I & J) and extended mouse survival compared to control treatment (p = 0.01; Cont-KO/control vs Con-KO/RT; Fig. 6K). Compared to control tumors, Abi-1 knockout significantly slowed tumor growth (Fig. 6I & J) and extended mouse survival (p = 0.02; Cont-KO/control vs Abi-1-KO/control; Fig. 6K), which could be further enhanced by radiation treatment (p = 0.03; Abi-1-KO/control vs Abi-1-KO/RT; Fig. 6K). Thus, Abi-1 mediates GBM resistance to genotoxic therapy *in vivo*.

### GTP Protects Normal Tissues from Genotoxicity through Rac1/Abi-1

In cancers such as GBM, the DDR mitigates the efficacy of genotoxic therapies. However, in non-transformed tissues, the DDR protects against genotoxic stressors encountered during DNA replication, cellular metabolism, and exposure to environmental agents, as well as the off-target effects of genotoxic cancer therapies on healthy tissues. To determine whether the GTP-Rac1-Abi-1 signaling axis mediates the DDR in normal tissues, we first irradiated murine enteroids with or without GTP supplementation. RT induced significant γ-H2AX positivity in enteroids, which could be significantly reduced by GTP supplementation (p < 0.0001; Fig. 7A & S7A). Consistent with our findings in GBM, RT induced a GTP-dependent dephosphorylation of Abi-1 in enteroids (Fig. 7B). We found a similar RT-induced and GTP-dependent dephosphorylation of Abi-1 in normal human astrocytes (Fig. 7C). Because astrocytes are more amenable to transfection than enteroids, we chose this model to confirm that Abi-1 dephosphorylation is dependent on Rac1 activity in normal tissues. Expression of constitutive active Rac1-Q61L promoted the dephosphorylation of Abi-1-S323 in normal human astrocyte after RT, while expression of dominant negative Rac1-T17N prevented Abi-1-S323 dephosphorylation (Fig. 7D), which is consistent with our findings in GBMs (Fig. 3). These data suggest that GTP promotes DNA repair and protects normal tissues from genotoxicity through Rac1/Abi-1 pathway.

**Figure 7.**
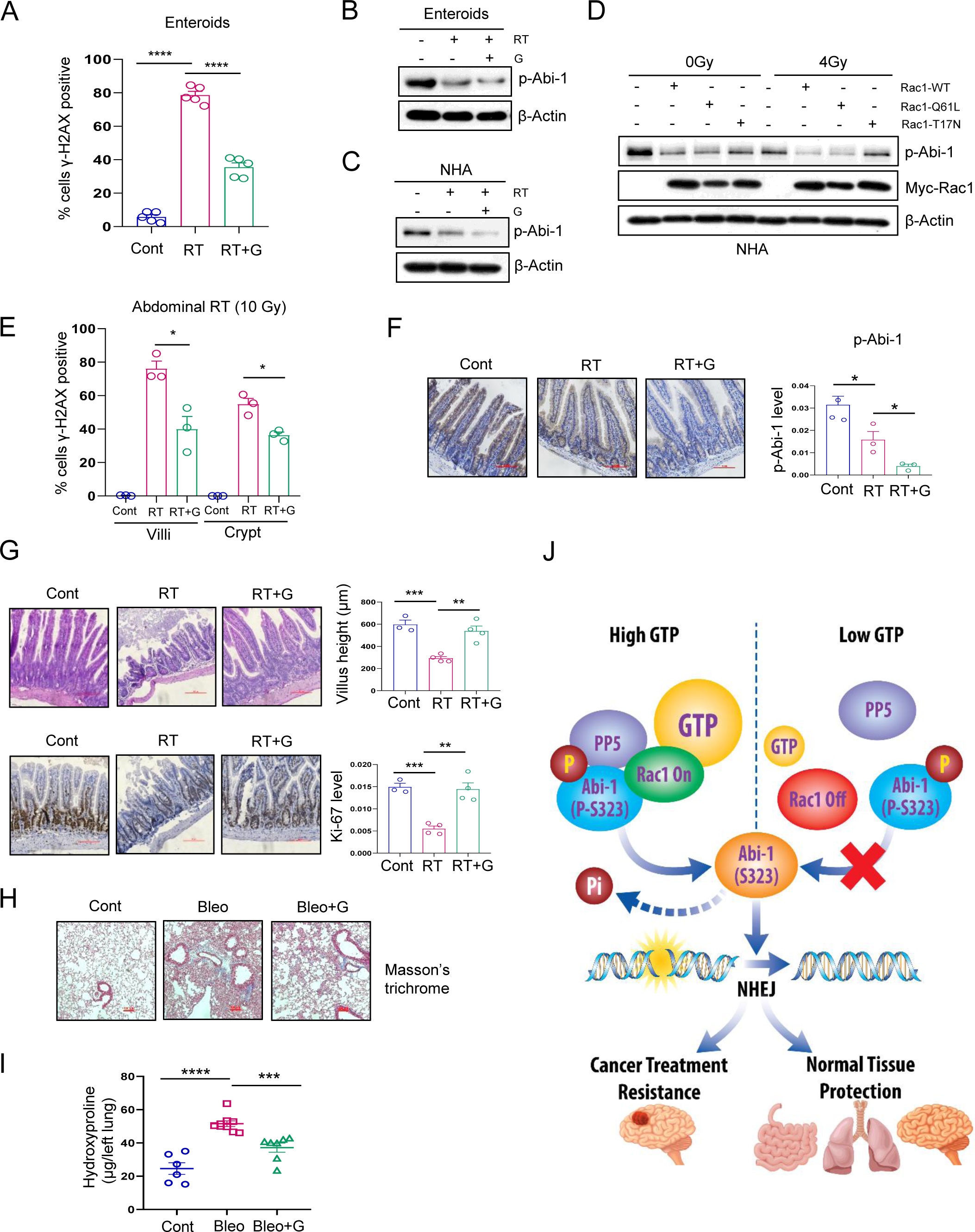
GTP protects normal tissues from genotoxicity through Rac1/Abi-1. (A) Enteroids were treated with RT (4 Gy) alone or RT (4 Gy) combined with G (50 μm) and harvested for IF staining 4 h post-RT. (B&C) Enteroids or normal human astrocytes (NHA) were treated as before and harvested for immunoblot assay. (D) NHA were transfected with wild type (WT), constitutively active (Q61L) or dominant negative (T17N) Rac1 plasmids and treated as above, and cells were harvested 4h post-RT for immunoblot. (E-G) C57BL/6J mice were treated with 7 doses of guanosine (300 mg/kg) by oral gavage or combined with one dose (10 Gy) of abdominal radiation. A subset of mice were sacrificed, and jejunums were harvested 4 h after receiving radiation for IF (E) or IHC staining (F), respectively. Another subset of mice continued to receive the rest of guanosine treatment and jejunums were harvested at day 14 for H&E (G, top panels) or ki-67 IHC staining (G, bottom panels). (H&I) Digital images of Masson’s trichrome staining for collagen deposition (blue) at ∼day 21 (H) and lung hydroxyproline content was quantified (I). (J) A schematic summary of our study.

We used two *in vivo* murine models to confirm our *in vitro* findings. We investigated RT-induced gastrointestinal injury and bleomycin-induced pulmonary fibrosis, which are common toxicities of genotoxic therapy encountered in the clinic. We performed both whole-body and abdominal radiation on C57BL/J6 mice and harvested jejunums for IF or IHC staining 4 hours post-RT. Both abdominal (Fig. 7E & S7B) and whole body (Fig. S7C & D) radiation caused significant DSBs on intestinal villi and crypts, which could be rescued by guanosine administration to increase GTP levels. Likewise, abdominal RT caused a dephosphorylation of Abi-1-S323 that was augmented by the administration of guanosine (Fig. 7F). To further interrogate RT-induced intestinal injury, we sacrificed and harvested jejunums at Day 14 after abdominal radiation, a time point at which RT-induced gut injury is apparent (34). RT caused villi shortening (Fig. 7G; top panels) and decreased crypt proliferation as measured by ki-67 staining (Fig. 7G; bottom panels), both of which were reversed by guanosine supplementation. We chose bleomycin-induced pulmonary fibrosis as an orthogonal model of genotoxic therapy-induced normal tissue injury (Fig. S7F). As expected, bleomycin-treated mice developed pulmonary fibrosis as assessed by Masson’s trichrome stain (Fig. 7H) and hydroxyproline assay (Fig. 7I). Supplementing GTP pools with guanosine treatment significantly ameliorated this bleomycin-induced fibrosis. Thus, we demonstrate that the GTP/Rac1/Abi-1 pathway can protect normal tissues from genotoxic stress.

## DISCUSSION

In this work, we have defined the molecular mechanism by which purines regulate the DDR and demonstrated its therapeutic relevance to different cell types and tissues. Surprisingly, purines promote DNA repair through GTP dependent signaling rather than by acting as physical substrates for DNA repair. High GTP levels activate the G protein Rac1, which stimulates PP5 to dephosphorylate Abi-1 at serine 323 (Fig. 7J). Dephosphorylated Abi-1 promotes double strand DNA break repair through non-homologous end joining. When Rac1-PP5-Abi-1 signaling is forcibly activated or inactivated, GTP pool sizes no longer regulates DNA repair. Disrupting this regulation using an inhibitor of Rac1 slows DNA repair in orthotopic GBM PDXs and improves the efficacy of radiation. GBM tumors and cell lines lacking Abi-1 or expressing an Abi-1 variant that cannot be dephosphorylated at serine 323 have increased sensitivity to genotoxic agents, but not other chemotherapies. This biology extends beyond GBM. GTP supplementation promotes Abi-1 dephosphorylation and DNA repair in normal astrocytes and enteroids and protects against normal tissue damage in mice treated with radiation or bleomycin. These findings implicate GTP as a key molecular link between metabolism and DNA repair and reveal numerous possibilities for therapeutic intervention.

Our work has implications for the treatment of GBM and suggests that a GTP-Rac1-PP5-Abi-1 signaling pathway mediates GBM resistance to standard of care genotoxic radiation and temozolomide. With this knowledge, we can design therapeutic strategies for GBM in which radiation and/or temozolomide are combined with inhibition of GTP-Rac1-PP5-Abi-1 signaling. Inhibition of GTP synthesis is the most actionable strategy to disrupt this signaling pathway in the clinic, as inhibitors of IMPDH, the rate limiting enzyme in GTP synthesis (both when it is formed de novo and from salvaged hypoxanthine), have been used in patients for decades (35,36). Inhibiting GTP synthesis also likely has a favorable therapeutic index in GBM. GBMs upregulate IMPDH expression and GTP synthesis to fuel nucleolar activity and tumorigenesis (37). The stem-like cells that are thought to mediate glioma initiation and recurrence also have heightened purine synthesis (38). We hypothesize that inhibition of IMPDH will selectively affect purine levels in glioma cells with high flux though this pathway and thus avoid potentiating the toxic effects of radiation and temozolomide on normal cortex, which has both low purine synthesis and low GTP demand. Our research team and others are currently conducting clinical trials combining the IMPDH inhibitor mycophenolate mofetil (MMF) with radiation and/or temozolomide for patients with GBM (NCT04477200, NCT05236036) (39). Such a strategy may also have promise in other cancers known to have elevated purine synthesis and IMPDH activity (40,41).

Inhibitors of the other members of this pathway (Rac1, PP5, Abi-1) are either in preclinical use (42) or non-existent. As these agents are developed and/or proceed to the clinic, it is possible that they could be useful in combination with radiation and/or temozolomide for the treatment of GBM. However, both Abi-1 and Rac1 are essential genes whose deletions are embryonic lethal (43,44). Given the numerous physiologic roles of Rac1 in cortical function (42) and the ubiquitous expression of Abi-1 in the cortex (45), it will be important to determine whether inhibition of these proteins potentiates the toxic effects of radiation and temozolomide on normal brain in addition to their therapeutic ability to overcome glioma treatment resistance.

GTP supplementation using the GTP precursor guanosine can protect normal tissues against genotoxic injury. These findings suggest that guanosine supplementation during a radiation disaster or space flight could be used as a toxicity-mitigation strategy. The practicalities of such an approach would depend on the timely administration of guanosine, as the NHEJ-mediated repair of RT-induced DNA damage is most critical within the first several hours of damage, with other repair pathways dominating later. Guanosine supplementation could also help mitigate the normal tissue toxicity of genotoxic cancer therapies. Boosting GTP levels in the rectum during radiation for prostate cancer or the lung during bleomycin treatment for Hodgkin’s lymphoma could diminish the toxicities associated with these treatments. It would be critical for such a strategy to avoid elevating GTP levels in cancer cells to avoid imparting treatment resistance, perhaps through formulations ensuring only localized delivery in the lung or GI tract.

There are many unanswered questions related to our findings. Why would cells evolve to preferentially rely on GTP signaling to stimulate DNA repair? Does the primacy for GTP in controlling DNA repair hold true even for cancers that are especially reliant on *de novo* pyrimidine synthesis for growth (46-48). Understanding this issue will determine how best to sequence and combine inhibitors of purine and pyrimidine synthesis with standard of care genotoxic agents for these cancers. How does the dephosphorylation of Abi-1 promote NHEJ? Can the currently available IMPDH inhibitors sufficiently penetrate the blood brain barrier to deplete GTP in GBM or are new drugs needed? And most importantly for our patients, can disrupting this biology safely improve outcomes in patients with GBM? The ongoing phase 0/1 clinical trials that we and others are conducting will begin to answer some of these questions, but careful laboratory study will be needed to fully understand this important new biology.

## METHODS

### Patient Samples

The GBM PDX samples used in Fig. 5A-F were obtained from The Mayo Clinic Brain Tumor Patient-Derived Xenograft National Resource (Overview - Translational Neuro-Oncology: Jann N. Sarkaria - Mayo Clinic Research). Treatment responses of dozens GBM PDX models were profiled previously (27,28). IHC assay (as shown below) was performed with indicated antibodies and correlation between protein expression and treatment benefit was analyzed by two-tailed unpaired student’s t test using Graphpad 8.0 software (Prism) (Fig. 5A-F). Relationship between Abi-1 expression and responses to treatment (TMZ alone or RT+TMZ) was assessed using Spearman correlation (Fig. S6B & C).

Human survival analysis of Rac1 and Rac1 pathway as shown in Fig. S5A & B was determined using the online tool KMplot (49). This is a web based, registration-free survival analysis tool that can perform univariate and multivariate survival analysis. Rac1 pathway gene signature was obtained from Gene Set Enrichment Analysis (BIOCARTA_RAC1_PATHWAY (gsea-msigdb.org) (as shown in Table S1) and used as an input for general overall survival analysis in GBM patients. Significance was computed using the Cox-Mantel log rank test.

### Cell Lines

HF2303 primary neurospheres, which were originally described by Dr. Tom Mikkelsen at Henry Ford Hospital (Detroit, MI), and RT-resistant GBM38 PDXs, which were obtained from Dr. Jann Sarkaria (The Mayo Clinic), were used as described in our previous study (11). Murine enteroids were a gift from Dr. Yatrik Shah (University of Michigan). Immortalized GBM adherent cell lines, DBTRG-05MG (DBTRG) and GB-1 were commercially purchased from American Type Culture Collection (ATCC) and Japanese Collection of Research Bioresources Cell BanK (JCRB), respectively. All cell lines were tested for mycoplasma positivity approximately monthly.

Immortalized DBTRG and GB-1 adherent cells and GBM38 PDX adherent cells were cultured in DMEM (Cat# 11965-092, Thermofisher Scientific) supplemented with 10% FBS (Cat# S11550, ATLANTA biologicals), 100 μg/mL Normocin (Cat# ant-nr-1, Invivogen) and 100 U/mL Penicillin-Streptomycin-Glutamine (Cat# 10378-016, Thermofisher Scientific). Primary patient-derived HF2303 neurospheres were cultured in DMEM-F12 (Cat# 11965-092, Thermofisher Scientific) supplemented with B-27 supplement (Cat# 17504-044, Thermofisher Scientific), N2 supplement (Cat# 17502-048, Thermofisher Scientific), 100 U/mL Penicillin-Streptomycin (Cat# 15140122, Thermofisher Scientific), 100 μg/mL Normocin, 20 ng/mL EGF (PeproTech, Cat# AF-100-15) and FGF (PeproTech, Cat# 100-18B). Enteroids were extracted from colonic crypts of C57BL/6J mouse (The Jackson Laboratory, RRID:IMSR_JAX:000664), plated in matrigel (Cat# 354230, Corning), and cultured in high Wnt containing media (L-WRN medium) provided by Michigan Medicine Translational Tissue Modeling Laboratory. Organoids were allowed to establish for at least 3-5 days as previously described (50).

### Animal Models

All mouse experiments were approved by the University Committee on Use and Care of Animals at the University of Michigan. 6-week-old male and female Rag1-KO mice (RRID:IMSR_JAX:002216), which were used for orthotopic tumor growth (Fig. 5, S5, and 6), and 6-week-old female C57BL/6J mice (RRID:IMSR_JAX:000664), which were used for enteroid establishment, intestinal-radiation model and bleomycin pulmonary fibrosis model (Fig. 7 & S7), were all obtained from the Jackson Laboratory.

### Constructs, siRNA, CRISPR/CAS9, and Transfection

pcDNA3.1(+)-3xFlag-tagged Abi-1 wild type and site-directed mutants (Abi-1-S323D and Abi-1-S323A) were commercially ordered from Sangon Biotech (http://www.life-biotech.com/; Shanghai, China) and mutation accuracy was confirmed by sequencing. pRK5-Myc-tagged Rac1 wild type (12985), Q61L (12983), and T17N (12984) plasmids were obtained from Addgene. SiRNAs of pan PP1 (a pool of 6 siRNAs, sc-43545), PP2A-Cα (a pool of 3 siRNAs, sc-43509), PP2A-Cβ (a pool of 3 siRNAs, sc-36301), PP4 (a pool of 3 siRNAs, sc-39202), PP5 (a pool of 3 siRNAs, sc-44602), Abi-1 (sc-417534-NIC) double nickase CRISPR/CAS9 knockout plasmids and control (sc-437281) were obtained from Santa Cruz Biotechnology. Puromycin (Cat# A1113803, Thermofisher Scientific) was used to select stable cell lines after control or Abi-1 knockout transfection. ON-TARGETplus SMART pool (a mixture of 4 siRNAs) of IMPDH1 (L-009687-00-0005), IMPDH2 (L-004330-00-0005), ADSS1 (L-009105-01-0005), and ADSS2 (L-009289-00-0005) were purchased from Dharmacon. Plasmid DNA or siRNA of phosphatases was transfected using Lipofectamine 2000 (Invitrogen; 52887) according to the manufacturer’s protocol. SMART pools from Dharmacon were transfected with DharmaFECT 1 Transfection Reagent (2001-01) according to the manufacturer’s protocol.

### Generation of the Phospho-Abi-1-S323 Antibody by Cell Signaling Technology

Rabbit polyclonal antibody was raised against phosphorylated Abi-1 (Ser323) by injecting rabbits with a synthetic, KLH-conjugated phosphopeptide corresponding to residues surrounding Ser323 of human Abi-1 protein as described previously (51). After being immunized subcutaneously with 0.5 mg of antigen in Complete Freund’s Adjuvant, rabbits were injected by five boosts of antigen (0.25 mg) in Incomplete Freund’s Adjuvant every three weeks thereafter. 10 days after the last antigen injection, crude bleeds were collected and antibody was purified using peptide affinity chromatography.

### Orthotopic Tumor Growth

GBM38 (Fig. 5G-N and Fig. 6I-K) and HF2303 (Fig. S5F-I) orthotopic mouse models were established as described previously (11). In brief, cells were infected with lentivirus harboring fluc (lenti-LEGO-Ig2-fluc-IRES-GFP-VSVG) and enriched for GFP-positive populations by flow cytometry. Luciferase-positive GBM38 (∼5×10^5^) and HF2303 (∼2×10^6^) cells were then orthotopically implanted in male and female Rag1-KO mice (RRID:IMSR_JAX:002216, The Jackson Laboratory). Brain tumor-bearing mice were randomized into four arms, including vehicle (Methy cellulose w/v 0.5%, Cat# B6385, Sigma-Aldrich, and tween 80 v/v 0.1%, Cat# P4780, Sigma-Aldrich, to 1x PBS), MBQ-167 alone, RT alone, or combined RT and MBQ-167 for Fig. 5 & S5, or two arms, including vehicle or RT alone for Fig. 6. MBQ-167 (75 mg/kg) and/or RT (2 Gy/fraction) were administered as shown in Fig. 5G and Fig. S5F, and RT (2 Gy/fraction) alone was administered as shown in Fig. S6P. For Fig. 5G, a subset of tumors analyzed for biologic endpoints (i.e. IHC) were taken 4 h after their second RT (or sham RT) dose. The rest of the animals (7-10 mice/group) continued to receive treatment and had tumor volume monitored. To assess the bioluminescence of brain tumors, mice were treated with 150 mg/kg D-luciferin (Cat# MB000102-R70170, Syd labs) by intraperitoneal injection and then imaged 10 min later using an IVIS™ Spectrum imaging system (PerkinElmer). Body weight and tumor volume were measured 1-2x weekly.

### Whole-body and Abdominal Radiation Mouse Models

For assessing radioprotection of GTP in normal intestinal tissues, 6-week-old female C57BL/6J mice (RRID:IMSR_JAX:000664, The Jackson Laboratory) were used and randomized into three arms (∼5 mice/arm), including saline vehicle, RT (10 Gy for both whole-body and abdominal model) alone, or combined RT and guanosine supplement (300 mg/kg by oral gavage) (Fig. S7E). Guanosine was administered 2 days before radiation and sustained for 7 days, and murine jejunum was harvested 4 h after whole-body (Fig. S7C & D) or abdominal radiation (Fig. 7E & F; S7B) to analyze p-Abi-1 levels and markers of DNA repair (IF or IHC staining) or 14 d after abdominal radiation for intestinal injury analysis (H&E and IHC staining; Fig. 7G).

### Bleomycin Pulmonary Fibrosis Mouse Model

To assess if GTP could protect normal lung tissues from bleomycin-induced fibrosis, 6-week-old female C57BL/6J mice (RRID:IMSR_JAX:000664, The Jackson Laboratory) were used and randomized into three arms with each experimental arm consisting of 10-15 mice. Treatments were performed as shown in Fig. S7F. Briefly, fibrosis was elicited in mice by oropharyngeal administration of a single dose of 1.5 units/kg body weight of bleomycin (Cat# B5507, Sigma-Aldrich); control mice received a volume of sterile saline equal to that described previously (52). Mice were sacrificed at the indicated time shown in Fig. S7F and lung tissues were harvested, with the left lung for hydroxyproline assay and the right lung lobes were assessed for histopathology as described below.

### Immunoblot Assay

GBM cells were transfected with siRNAs, DNA plasmids, and/or treated with individual compounds or radiation, and harvested at indicated time points. Cells were lysed using RIPA lysis buffer (Cat# 89900, Thermofisher Scientific) supplemented with PhosSTOP phosphatase inhibitor (Cat# 04906845001, Roche) and complete protease inhibitor tablets (Cat# 1187358001, Roche). Protein expression was detected with individual antibodies.

### Rac1 Activity Assay

Rac1 activity assay was performed according to the manufacturer’s protocol (Cat# BK035, Cytoskeleton Inc). DBTRG or HF2303 cells were transfected with Rac1-WT, -Q61L, and -T17N plasmids, and treated with MPA, guanosine, and/or radiation. Cells were harvested 4 h post-RT and lysed using lysis buffer with protease inhibitor cocktail (all included in the same kit). ∼500 μg protein for each group were incubated with PAK-PBD beads (10-20 μl) at 4 ^0^C for 1 h, followed by bead washing with kit-included washing buffer and eluting with 20 μl 2x Laemmli sample buffer (Cat# 161-0737, BIO-RAD). A 20-50 μg sample for each group was saved for immunoblot quantitation of total Rac1. Kit-included pure Rac1 protein, GTPγS (active control), or GDP (inactive control) were used based on the manufacturer’s protocol.

### Immunofluorescence Assay

Immunofluorescence (IF) was performed as described in our previous study (11). Briefly, immortalized or primary GBM cells were plated and treated with indicated conditions. γ-H2AX foci were detected with mouse monoclonal antibody anti-phospho-Histone H2AX (Ser139) (1:1000 dilution; Cat #05-636, Millipore) and goat anti-mouse IgG secondary antibody, alexa fluor 594 (1:1000 dilution; Cat# A-11005, Thermofisher Scientific) in immortalized cells and neurospheres, and with rabbit anti-phospho-Histone H2AX (Ser139) (1:100 dilution; Cat #9718, Cell Signaling Technology) and goat anti-rabbit IgG secondary antibody, alexa fluor 594 (1:1000 dilution; Cat# ab150080, Abcam) in enteroids and mouse intestinal tissues. Rad51 foci were detected with anti-mouse primary antibody (1:300 dilution, Cat# GTX70230, GeneTex) and anti-mouse IgG secondary antibody, alexa fluor 594 as described above. γ-H2AX foci were evaluated with ImageJ (FIJI) software and scored for each condition in ∼100 cells of immortalized GBM cells, ∼10 spheres of primary GBM cells, 10-15 enteroids, and ∼30 villi and ∼90 crypts for each mouse intestinal tissues. The foci threshold of γ-H2AX and Rad51 foci was 10 for immortalized cells and intestinal tissues and 3-5 for primary GBM neurospheres and enteroids.

### Celltiter-Glo Cell Viability Assay

HF2303 primary or DBTR immortalized GBM cells with control or Abi-1-KO, or re-expression of Abi-1-WT, -S323D or -S323A, were plated into 96 well plates and treated with different doses of radiation or escalating doses of indicated compounds as shown in Fig. 6 & S6. After growing for 3 days (for non-genotoxic paclitaxel and vincristine) or 7 days (for radiation and other genotoxic compounds), cell viability was detected using CellTiter-Glo 2.0 (Cat# G9241, Promega) in DBTRG and CellTiter-Glo® 3D (Cat# G9681, Promega) in HF2303 following the manufacturer’s protocol.

### H&E and Immunohistochemical Assay

Mouse tumors, jejunum, or lung tissues, were harvested, fixed in 10% formalin, and embedded in paraffin for hematoxylin & eosin (H&E) or immunohistochemical (IHC) staining. IHC assay was performed using the ABC Vectastain Kit (Cat# PK6101 or PK6102, Vector Laboratories) as described previously (11). After deparaffinization, rehydration, antigen retrieval, and blocking, tumor or normal organ tissue slides were incubated with primary antibody of Ki-67 (1:10000 dilution; Cat# 550609, BD Biosciences), p-Abi-1-S323 (1:500 dilution, this paper), p-Pak1/2(Thr423/402) (1:300 dilution; Cat# 2601, Cell Signaling Technology), p-Pak2 (1:300 dilution; Cat# 2607, Cell Signaling Technology), p-γ-H2AX (1:1000 dilution; Cat# 9718, Cell Signaling Technology) at 4 °C overnight. After being incubated with the secondary antibody for 30 min, the tissue slides were stained with DAB substrate kit (Cat# SK-4100, Vector Laboratories) and finally counterstained with hematoxylin, dehydrated and mounted. IHC quantification of each protein was performed as described previously (53). Villus heigh of jejunum was measured using ImageJ software (FIJI).

### Clonogenic Survival Assay

DBTRG cells with Abi-1-KO or Abi-1-WT, -S323D and -S323A re-expression were irradiated with indicated doses, following by replating at clonal density. Plates were stained and colonies containing > 50 cells were counted after 10 to 14 days of growth. The RT enhancement ratio (ER) was calculated as the ratio of the mean inactivation dose (Dmid) under control conditions (Cont-KO or Abi-1-WT) divided by the mean inactivation dose under Abi-1 knockout or Abi-1-S323D and -S323A overexpression treatment.

### NHEJ Linearized pEYFP Plasmid Assay (qPCR)

pEYFP-N1, which was originally obtained from Clontech Laboratories, was a gift from Dr. Meredith Morgan (University of Michigan). Plasmid was linearized by digestion with NheI enzyme between the promoter and coding sequence of EYFP, followed by gel-purification with PCR purification Kit (Qiaquick CAT#28104) for the linear products. GBM cells were treated with individual purine supplements (A or G), MPA (10 μm), or AG2037 (150 nm) overnight (Fig. 1), or first transfected with individual Abi-1 plasmids and then treated with guanosine supplement, MPA (10 μm), or combined MPA and G (Fig. 2), and cells were all retreated with purine supplements 2 h before transfection of linearized pEYFP products (0.5 μg/well for 12-well plate), followed by cell harvesting (24 h post-transfection) and DNA extraction with QIAprep Spin Miniprep kit (Cat# 27106). DNAPK inhibitor M3814 (Selleckchem, Cat# S8586) (54), which inhibits NHEJ repair, was used as a positive control. The efficiency of end-joining DNA repair ability was assessed by SYBR green (Cat# 4385612, Thermofisher Scientific) real time qPCR of the rejoined EYFP region, which was normalized to an uncut flanking DNA sequence, relative to DMSO-treated (Fig. 1) or Abi-1-WT (DMSO-treated) cells (Fig. 2).

### NHEJ and HR I-Scel Reporter Assay (Flow cytometry)

The I-SceI-based NHEJ or HR assay was performed as described previously (55). Immortalized GBM cells or primary neurospheres stably expressing NHEJ or HR reporter were seeded in 6-well plates and treated with adenosine or guanosine supplements overnight and retreated with each purine supplement and/or MPA (10 μm), AG2037 (150 nm) 2 h before being infected by I-Scel adenovirus. DNAPK inhibitor M3814 (Selleckchem, Cat# S8586) (54) and ATM kinase inhibitor Ku-60019 (Cat# 17502, Caymanchemical) (56), was used as NHEJ and HR positive control, respectively. Cells were harvested after 48 h and the percentage of GFP-positive cells (indicative of NHEJ or HR repair) was quantified by flow cytometry.

### Hydroxyproline Assay

Hydroxyproline content of the lung was measured as previously described with modifications (57). Lungs were perfused, harvested, and homogenized in 1ml of PBS with protease inhibitor with EDTA (Cat# 11836153001, Roche). 1 ml 12N HCL was then added to the homogenate and samples were hydrolyzed at 120°C for 24 hours. Thereafter, 5 μl of each sample or standard was combined with 5 μl citrate-acetate buffer (5% citric acid, 1.2% glacial acetic acid, 7.24% sodium acetate, 3.4% sodium hydroxide into 100ml ddH2O) in a 96-well plate. 100 μl of chloramine T solution (0.282 g chloramine T to 16 ml of citrate-acetate buffer, 2.0 mL of N-propanol, and 2.0 ml ddH2O) was added and samples incubated for 30 minutes at room temperature. Next, 100 μl of Ehrlich’s reagent (2.5 g 4-dimethylaminobenzaldehyde added to 9.3 ml of N-propanol and 3.9 ml of 70% perchloric acid) was added, and the samples were incubated at 70°C for 30 minutes. The absorbance of each sample was then measured at 550 nm. Standard curves for the experiment were generated using cis-4-Hydroxy-1-proline (Sigma) with stock solution 4 mg/mL and sequential dilution range from 400 ug/ml down to 1.5625 ug/ml.

### Masson’s Trichrome Histopathology Assay

Lungs were fixed in 10% formalin and embedded in paraffin. Sections with 5-μm thick were stained with NovaUltra Masson Trichrome Stain Kit (Cat# IW-3006, IHCWORLD) according to the manufacturer’s protocol to detect collagen and visualize the extent of fibrosis.

### Phosphoproteomic Assay

The phosphoproteomic assay shown in Fig. 2A was performed by PTM BIOLABS (https://www.ptmbiolabs.com/). Briefly, HF2303 GBM neuroshperes were dissociated and plated. 3-4 days later, formed neurospheres were treated with DMSO control, MPA (10 μm), and/or guanosine supplement (50 μm) overnight, and retreated with guanosine (50 μm) 2 h before RT (6 Gy), followed by cell harvesting 4 h post-RT. The abundance of phosphopeptides was determined by PTM BIOLABS as described previously (58,59). Data from this profiling effort are attached as a separate supplementary file (Supplementary Data 1).

### Expression and Treatment Resistance Analyses

Coessentiality of Rac1 and Abi-1 was determined using DepMap (60). Briefly, gene effect scores were calculated using Chronos where a negative score indicates growth inhibition or death upon CRISPR knockout and correlation was assessed with a Spearman’s correlation test. Correlation of Abi1 transcript expression and bleomycin sensitivity (AUC) was also performed using DepMap across 32 glioma cell lines. Correlation of Abi-1 transcript expression and GBM PDX treatment responses was performed as we have done previously (28).

### Statistical Analysis

γ-H2AX or Rad51 foci formation, pEYFP NHEJ activity, I-Scel-NHEJ or HR activity, clonogenic survival, IC50 of individual compound, hydroxyproline level, and PDX or mouse tumor IHC staining after transfection and/or compound treatment were analyzed by unpaired two-tailed t-tests using GraphPad Prism Version. Tumor volume of orthotopic GBM was normalized to one at the first day for each group. Mouse survival was estimated by the Kaplan– Meier method and compared using the log-rank (Mantel-Cox) test. Significance threshold was set at p < 0.05.

## Authors’ Disclosures

S.G.Z reports unrelated patents licensed to Veracyte, and that a family member is an employee of Artera and holds stock in Exact Sciences. M.A.M gets research support and honoraria from AstraZeneca. No disclosures were reported by the other authors.

## Author Contributions

D.R.W., and W.Z. conceptualized and designed the study. W.Z. and D.R.W led and developed the methods. W.Z., Z.Z., A.L., J.Y., J.X., A.Y., and Y.Y. performed the experiments. W.R.K., J.L., and J.J. provided material support. J.L. and A.W.W. generated p-Abi-1 antibodies. S.S. helped and Y.S. supervised the experiments of gastrointestinal models. J.S. and N.W. helped, and M.P.G. supervised the bleomycin animal model. A.J.S., A.U.K., E.R.P., N.K., C.K.W., S.P., and A.M.O. helped and coordinated with the animal experiments. J.N.S. supported PDXs samples and cells. S.G.Z. and B.G. provided bioinformatic support. A.L.E., S.P.F., S.Z., W.A.H., Y.U., M.A.M., T.S.L., C.A.L., and Y.S reviewed the paper and provided material support. W.Z. and D.R.W. supervised the study, analyzed, and interpreted the data, wrote, reviewed, and revised the paper.

## Acknowledgments

We thank Steven Krongenberg for his assistance with illustrations. D.R.W. was supported by grants from the Forbes Institute for Cancer Discovery, the NCI (K08CA234416; R37CA258346), the NINDS (R01NS129123), the Damon Runyon Cancer Foundation, the Sontag Foundation, the Ivy Glioblastoma Foundation, Alex’s Lemonade Stand Foundation, and the Chad Tough Defeat DIPG foundation. S.S was supported by a Crohn’s and Colitis Foundation Research fellow award (623914). A.J.S was supported by NCI (F32CA260735). S.G.Z was supported by DP2 OD030734. B.G was supported by the National Research, Development and Innovation Office RRF-2.3.1-21-2022-00015 and TKP2021-NVA-15. M.A.M was supported by NCI (R01CA240515) and UMCCC Core Grant (P30CA046592). M.P.G was supported by NIH (R35HL144979). C.A.L. was supported by a 2017 AACR NextGen Grant for Transformative Cancer Research (17-20-01-LYSS) and an ACS Research Scholar Grant (RSG-18-186-01). Y.M.S was supported by NCI (R01CA148828, R01CA245546, and R01DK095201) and UMCCC Core Grant (P30CA046592).

