## Supplemental Information for "GTP signaling links metabolism, DNA repair, and responses to genotoxic stress"

**Running title:** GTP signaling links DNA repair to genotoxic therapies

**SUPPLEMENTAL INFORMATION**

**FIGURE LEGENDS**

**Figure S1 GTP promotes DSB repair through NHEJ.**

(A&B) Representative images of γ-H2AX (S139) foci staining in HF2303 (corresponding to Fig. 1A) and DBTRG (corresponding to Fig. 1B). (C&D) GB-1 cells were treated with adenosine (A; 50 μm), guanosine (G; 50 μm) or pooled purine and pyrimidine nucleosides (Nuc; 8x) for 24 h and retreated with the same doses of individual purine or pooled nucleosides 2 h before RT and harvested for γ-H2AX quantification by IF 4 h post-RT. Representative images of γ-H2AX (S139) foci were shown (D). (E&F) Key enzymes of GTP or ATP synthesis were silenced by a SMARTpool of siRNAs and silencing efficiency was tested by immunoblot with indicated antibodies (E), or cells were treated with individual purine supplement post transfection and RT sensitivity was evaluated by γ-H2AX foci IF staining (F). (G) Cells were treated with AG2037 (150 nm) alone or combination with purine supplements (A or G and then harvested 4 h post-RT for γ-H2AX quantification by IF. (H) Cells expressing the I-SceI-HR reporter were treated with A (50 μm), G (50 μm), the GTP-only inhibitor mycophenolic acid (MPA, 10 μm), the dual GTP/ATP inhibitor AG2037 (150 nm) or their combinations. DNA damage was induced by infection with I-SceI adenovirus and HR was assessed by quantifying the percentage of cells GFP positive 48 h later and normalized to untreated cells. The ATM kinase inhibitor Ku-60019 was used as a positive control (HR-in). (I&J) Cells were treated with MPA (10 μm) or G (50 μm) alone or in combination, and retreated with G 2 h before RT, followed by Rad51 foci IF staining 4 h after radiation. Data are presented as mean ± SEM from three biologically independent experiments. Two-tailed unpaired student's t test *p < 0.05, **p < 0.01, ***p < 0.001, ****p < 0.0001.

**Figure S2 Dephosphorylation of Abi-1-S323 is crucial to GTP-induced NHEJ.**

(A) Plasmids encoding Abi-1 wildtype (WT) and site-directed mutants (S323D phospho-mimetic mutant and S323A dephospho-mimetic mutant) were established to test the specificity of newly-generated p-Abi-1-S323 antibody. Abi-1-KO cells were transfected with Abi-1-WT, -S323D, and -S323A plasmids and immunoblot was performed to detect protein expression. (B) Cells were treated with MPA (10 μm) and/or G (50 μm) for 24 h and retreated with G (50 μm) 2 h before RT, followed by cell harvesting 4 h after RT for immunoblot assay. (C&D) Control knockout (Cont-KO) or Abi-1 knockout (Abi-1-KO) cells were treated with MPA, G, and/or RT (4 Gy) and harvested as above to assess γ-H2AX levels by immunoblot (C) or foci quantification (D). (E&F) Cells expressing wild type (WT), phosphomimetic (S323D) or dephosphomimetic (S323A) Abi-1 were treated with RT (4 Gy), MPA (10 μm) and/or G (50 μm) and analyzed for γ-H2AX levels by immunoblot (E) or foci quantification by IF (F). (G&H) Abi-1-KO cells were transfected with individual Abi-1 plasmids and treated as in S2E & S2F, followed by Rad51 foci IF staining. Data are presented as mean ± SEM from three biologically independent experiments for Fig. D&H and two biologically independent experiments for Fig. F&G. Fig. A, B, C, E are representative figures from three biologically independent experiment. Two-tailed unpaired student's t test *$p < 0.05$, **$p < 0.01$, ***$p < 0.001$, ****$p < 0.0001$.

**Figure S3 Rac1 controls the GTP-dependent dephosphorylation of Abi-1-S323.**

(A) DBTRG cells were transfected with individual Rac1 plasmids and Rac1 activity assay was performed according to manufacturer's protocol. In brief, cell pellets were lysed and equal amount protein solution was incubated with GST-tagged PAK-PBD beads, which specifically binds to GTP-bound Rac1. One hour later, beads were washed and eluted with 2 x loading buffer for immunoblot. Purified Rac1 protein was used as a positive control, as was GTPγS-treatment of cell lysates. GDP treatment of cell lysates was used as a negative control. (B&C) Cells were transfected with plasmids encoding Rac1-WT, -T17N (dominant negative), or -Q61L (constitutively active), and treated with MPA, G, or RT as in Figure 2. Cells were harvested 4h post RT (4 Gy) to assess Rac1 activity (B) or γ-H2AX foci by immunofluorescence (C). In (B), Long E. indicates long exposure of film and Short E. indicates short exposure. (D&E) Cont-KO or Abi-1-KO cells were transfected with constitutively active Rac1-Q61L and then irradiated (4 Gy), followed by immunoblot (D) or γ-H2AX foci IF staining (E). Data are presented as mean ± SEM from three biologically independent experiments for Figure C & E and Figure B & D are representative figures from three biologically independent experiment. Control data in Figure A is from a single experiment. Two-tailed unpaired student's t test *$p < 0.05$, **$p < 0.01$, ***$p < 0.001$, ****$p < 0.0001$.

**Figure S4 Protein phosphatase 5 mediates the GTP/Rac1-depdendent dephosphorylation of Abi-1 (S323) and downstream DSB repair.**

(A&B) Cells were treated with the phosphatase inhibitors okadaic acid (OA; 15 nm) or fostricein (Fos; 100 nm) 1 h before RT (A), or first transfected with pan PP1 siRNA, mixture of PP2A catalytic (PP2A-C) subunit α/β, PP4 and PP5 for 48 h and irradiated after transfection (B), followed by cell harvesting 4 h post-RT for immunoblot. (C-F) Cells were transfected with constitutively active Rac1-Q61L and then treated with okadaic acid (15 nm) or fostricein (100 nm) 1 h before RT (C&D), or transfected with individual phosphatase siRNA pool, along with overexpression of Rac1-Q61L or control (E&F), followed by immunoblot (C&E) or γ-H2AX foci IF staining (D&F). Data are presented as mean ± SEM from three biologically independent experiments for Figure D, F and Figure A, B, C, E are representative figures from three biologically independent experiments. Two-tailed unpaired student's t test $*p < 0.05$, $**p < 0.01$, $***p < 0.001$, $****p < 0.0001$.

**Figure S5 Rac1 activity influences GBM treatment responses.**

(A&B) Correlation between expression of Rac1 (A) and Rac1 pathway genes (B) and patient survival were analyzed and plotted using KM plotter. (C) Rac1 and Abi-1 gene essentiality scores were obtained from Depmap. Glioma cell lines are indicated with a star. Gene effect scores were calculated using Chronos. (D) DBTRG or GBM38 stable cells with control or Abi-1 knockout were injected into mouse flank (DBTRG) or brain (GBM38). Tumors were harvested and fixed with 10% formalin for IHC staining with newly generated p-Abi-1 S323 antibody. (E) Representative IHC pictures of p-Abi-1-S323, p-Pak1/2 (Thr423/402), p-Pak2 (Ser20), and γ-H2AX (corresponding to Fig. 5H-K). (F) A schematic timeline of HF2303 orthotopic mouse models. (G-I) Luciferase-positive HF2303 patient-derived neurospheres were implanted orthotopically and mice were treated with Rac1 inhibitor and/or RT according to the timeline shown in Fig. S5F. Bioluminescence imaging was performed 10 min after injection of 150 mg/kg D-luciferin (G). Total flux of equal-area ROIs at each time point were normalized to flux at the first day of treatment for evaluating tumor progression (H). Mice were monitored and Kaplan–Meier survival curve was plotted (I). Data are presented as mean ± SEM from 5-6 independent mice for Fig. S5G-I. Log-rank (Mantel-Cox) test $*p < 0.05$, for Fig. S5I.

**Figure S6 Abi-1 mediates genotoxic treatment efficacy in GBM.**

(A) Gene expression and bleomycin sensitivity were obtained from and plotted using Depmap. (B&C) Cont-KO or Abi-1-KO cells were irradiated with different doses (D) or treated with escalating concentrations of bleomycin (E; Cont-KO vs Abi-KO IC50:24.11 ± 8.27 vs 0.96 ± 0.32, p = 0.049), TMZ alone (F; 128.25 ± 6.16 vs 74.60 ± 7.27, p=0.044) or TMZ combined with RT (4Gy) (G; 135.20 ± 25.39 vs 64.59 ± 2.12, p = 0.045) and cell viability were evaluated with clonogenic assay (D) or long-term cell viability assay (E-G). (H-K) Cells were treated with escalating concentrations of paclitaxel (H&I; DBTRG IC50: 0.007 ± 0.0001 vs 0.009 ± 0.001, p = 0.068; HF2303 IC50: 0.029 ± 0.015 vs 0.018 ± 0.01, p = 0.565) or vincristine (J&K; DBTRG IC50: 0.033 ± 0.004 vs 0.064 ± 0.029, p = 0.366; HF2303 IC50: 0.018 ± 0.007 vs 0.095 ± 0.070, p = 0.3339), and cell viability was evaluated 3 days after treatment. (L-O) Abi-KO DBTRG cells were transiently transfected with Abi-1-WT, -S323D or -S323A, followed by irradiation (L), bleomycin (M; IC50 of WT vs S323D:0.111 ± 0.004 vs 0.035 ± 0.008, p = 0.021;  WT vs S323A: 0.111 ± 0.004 vs 0.160±0.007, p = 0.038), TMZ alone (N; IC50 of WT vs S323D: 232.35 ± 14.49 vs 92.92 ± 8.80, p = 0.02;  WT vs S323A: 232.35 ± 14.49 vs 466.20 ± 40.91, p = 0.048) or TMZ combined with RT (4Gy) (O; IC50 of WT vs S323D:174.35 ± 15.47 vs 9.92 ± 4.84, p = 0.014; WT vs S323A: 174.35 ± 15.47 vs 256.75 ± 0.78, p = 0.049) treatment as discussed above. All drug concentrations are in μM. (P) A schematic timeline of GBM38/Abi-1-KO orthotopic mouse models. Figures D to O are representative figures from two to three biologically independent experiments (mean ± SEM).

**Figure S7 GTP protects normal tissues from genotoxicity through Rac1/Abi-1.**

(A&B) Representative images of γ-H2AX (S139) foci staining in enteroids (corresponding to Fig. 7A) and small intestine tissues of abdominal radiation mouse model (corresponding to Fig. 7E). (C&D) C57BL/6J mice were treated with 7 doses of guanosine (300 mg/kg) by oral gavage and/or one dose (10 Gy) of whole-body radiation. Mice were sacrificed and jejunums were harvested 4 h after receiving radiation for IF staining. (E) A schematic timeline of abdominal or whole-body radiation mouse models. (F) A schematic timeline of bleomycin-induced fibrosis mouse models. C57BL/6J mice were treated with 10 doses of guanosine (300 mg/kg) by oral gavage and/or a single oropharyngeal dose of 1.5 units/kg body weight. Mice were sacrificed and lung tissues were harvested for Masson’s trichrome staining or hydroxyproline assay.

#### Table S1 Rac1 Signaling Pathway

| Gene name | Correlation with Rac1 | Reference |
|---|---|---|
| PIK3R1 (P85α) | Positive | [1-4] |
| PIK3CA (P110α) | Positive | [4, 5] |
| NCF2 | Positive | [6-11] |
| WASF1 (WAVE) | Positive | [12] |
| CHN1 | Negative | [13-15] |
| PAK1 | Positive | [16-18] |
| PLD1 | Positive | [19] |
| CDK5 | Positive | [20] |
| LIMK1 | Positive | [21, 22] |
| ARFIP2 | Positive | [23-25] |
| VAV1 | Positive | [26] |
| PIK3CG (P110γ) | Positive | [4] |
| RPS6KB1 | Positive | [27,28] |
| CDK5R1 | Positive | [29, 30] |
| MAP3K1 | Positive | [31] |
| RALBP1 | Negative | [32] |
| TRIO | Positive | [33, 34] |
| RAC1 | Positive | |

### Zhou etal., Figure S1

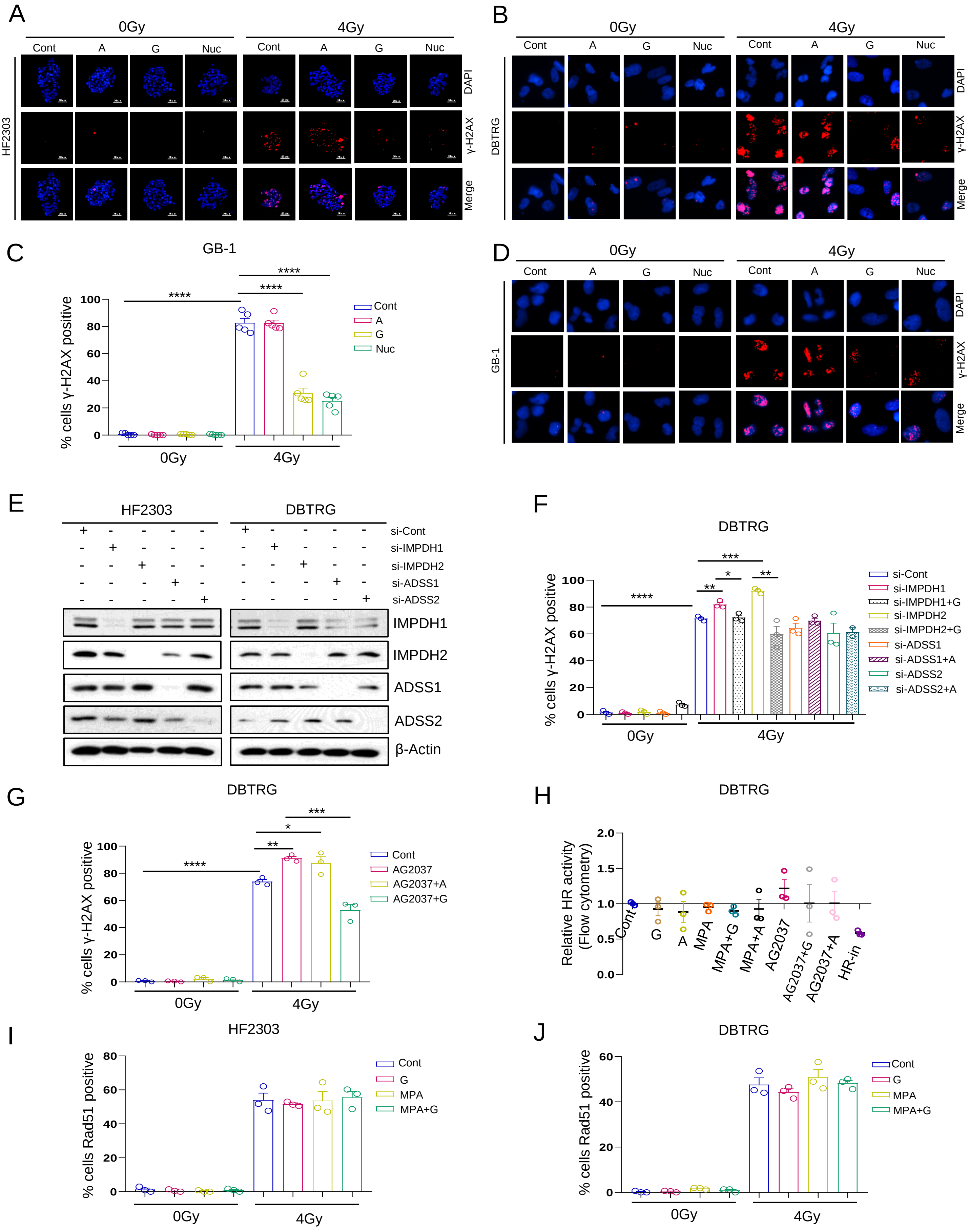

*Zhou etal.*, Figure S2

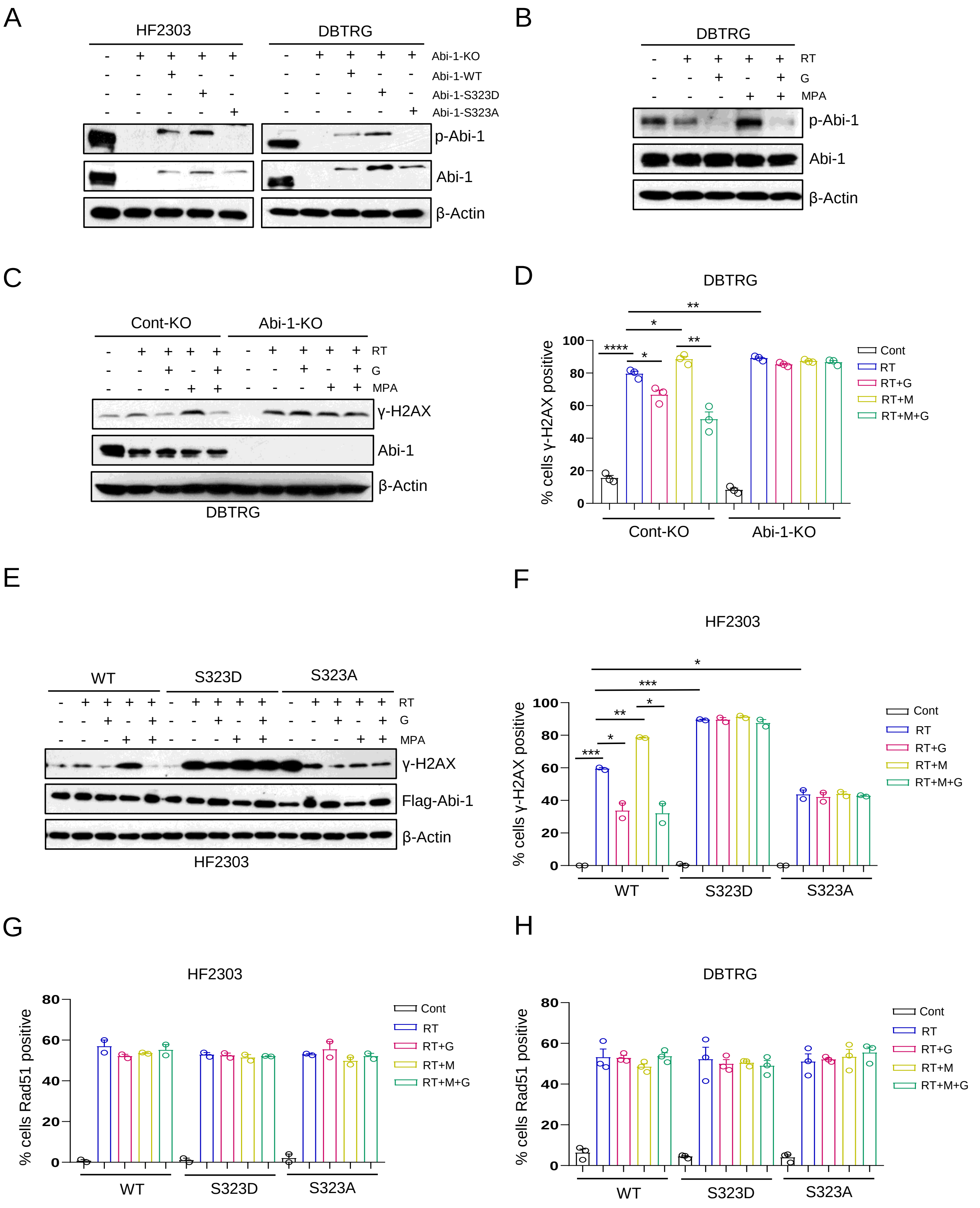

*Zhou etal.*, Figure S3

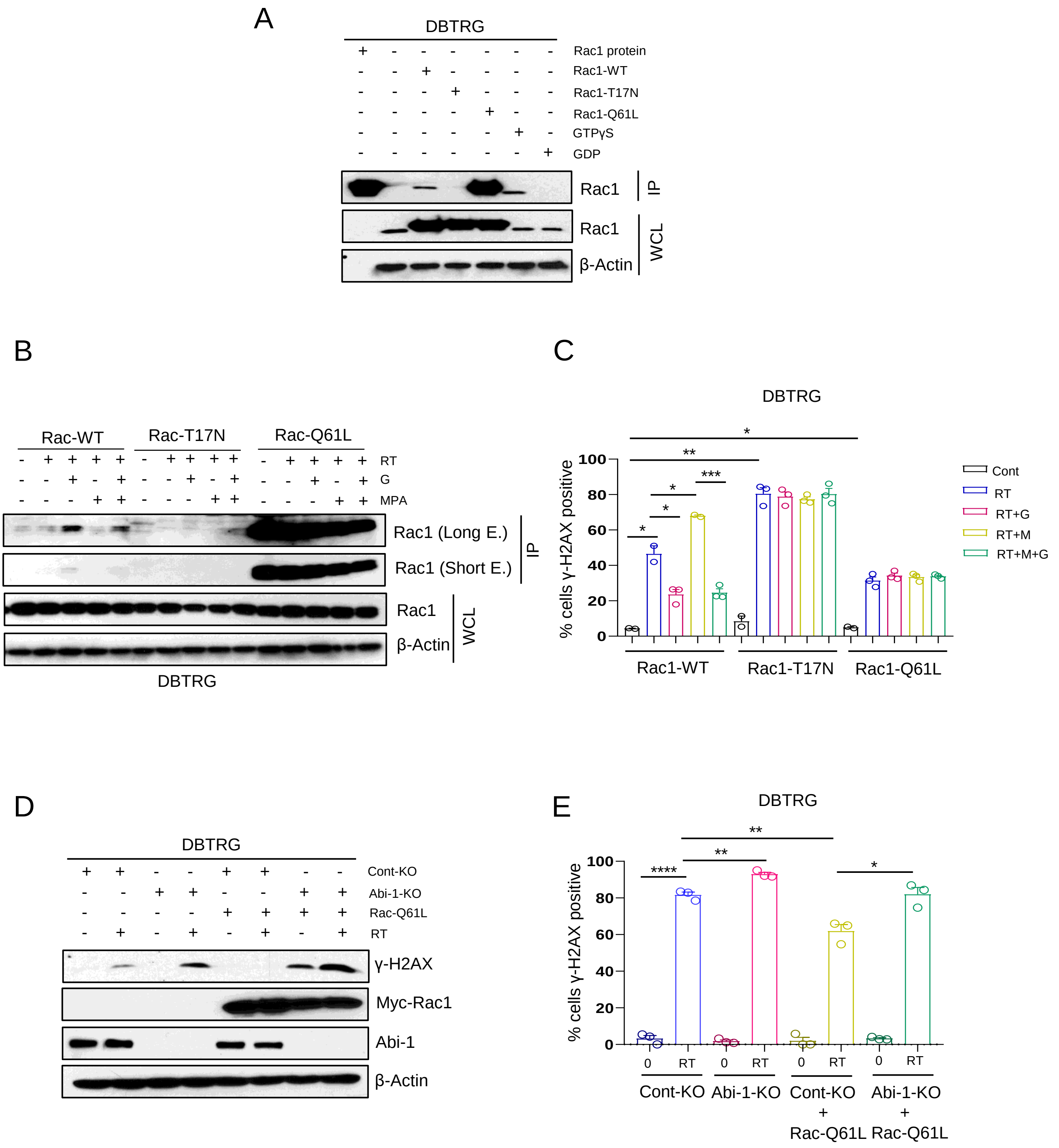

*Zhou etal.*, Figure S4

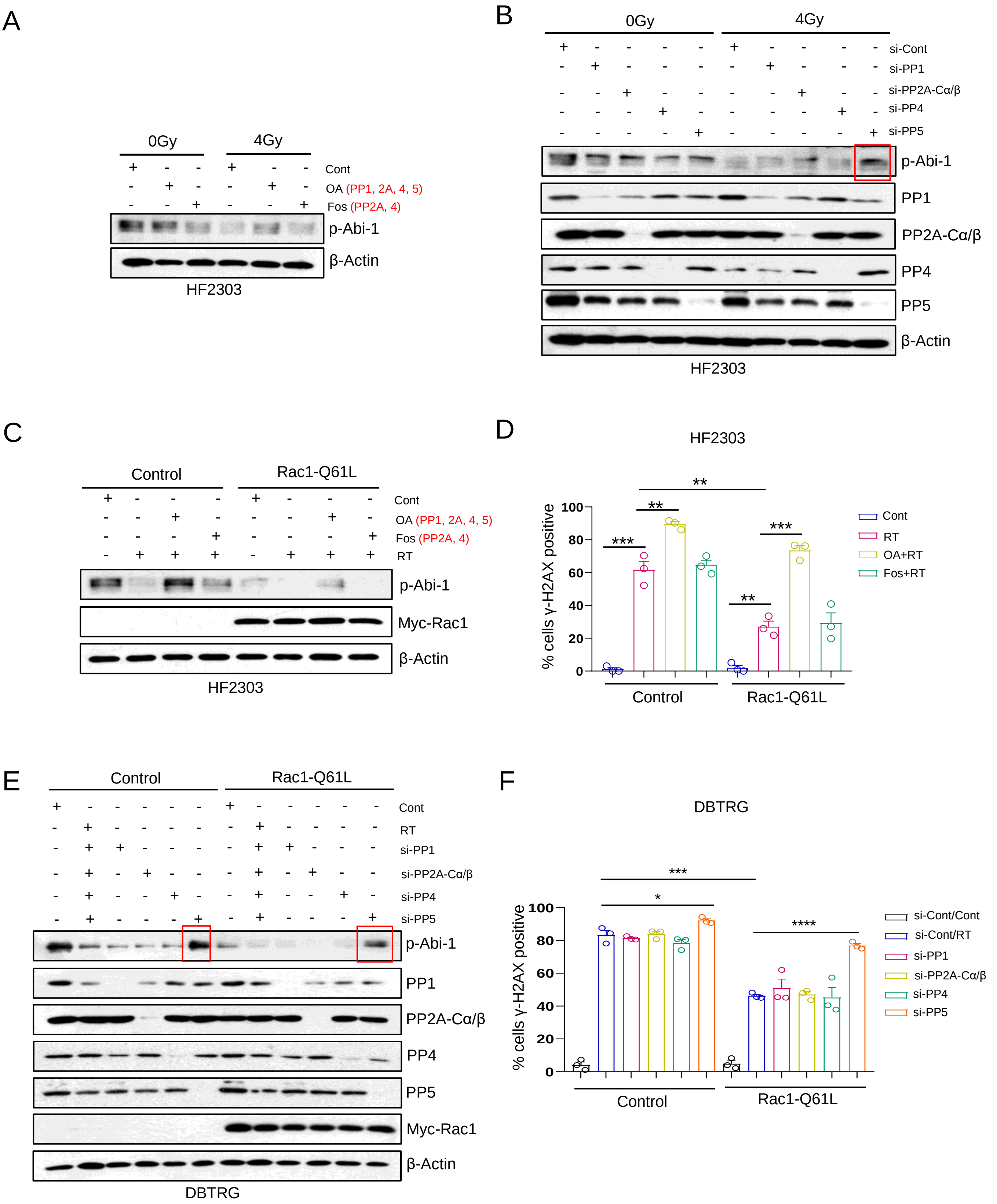

*Zhou etal.*, Figure S5

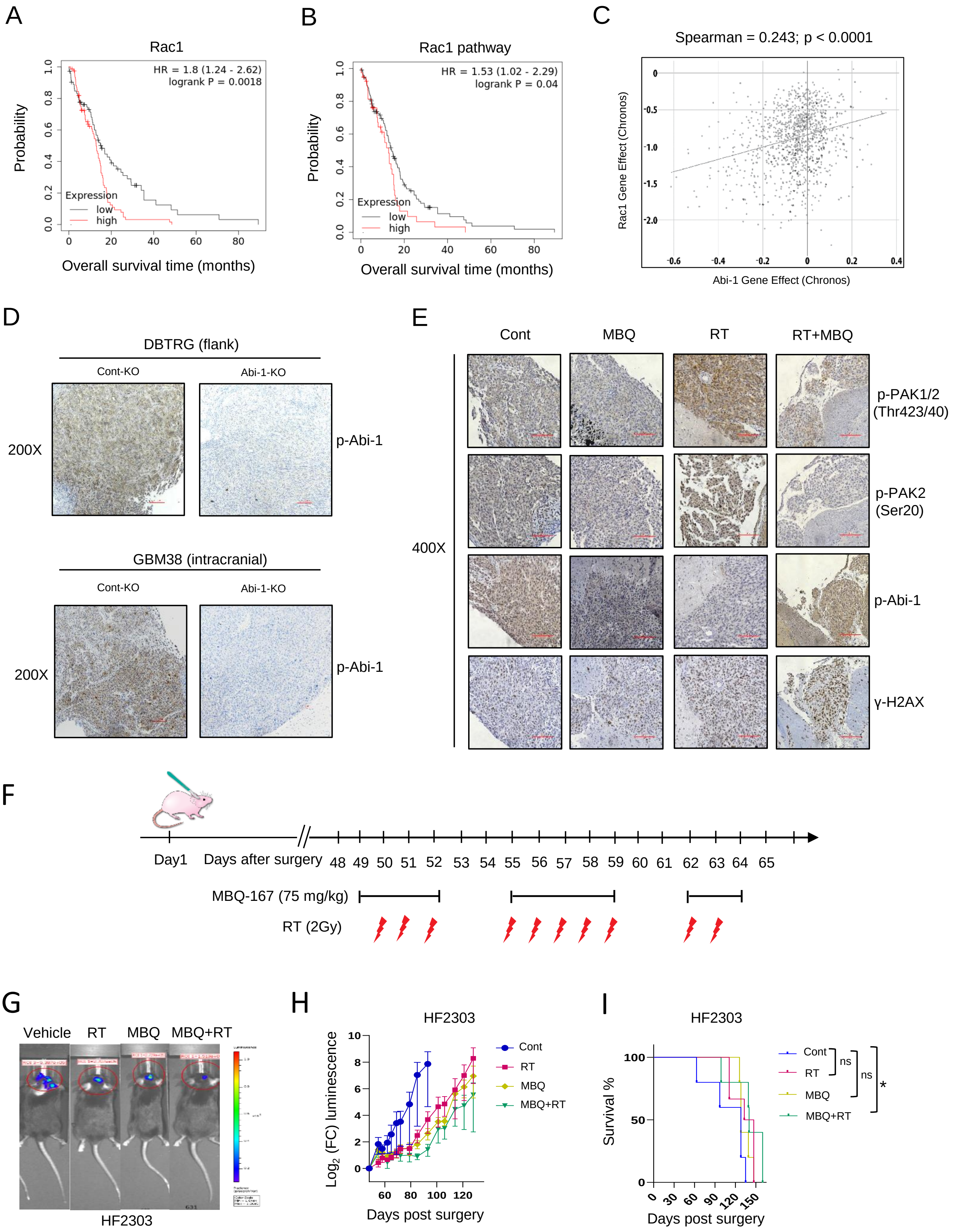

*Zhou etal.*, Figure S6

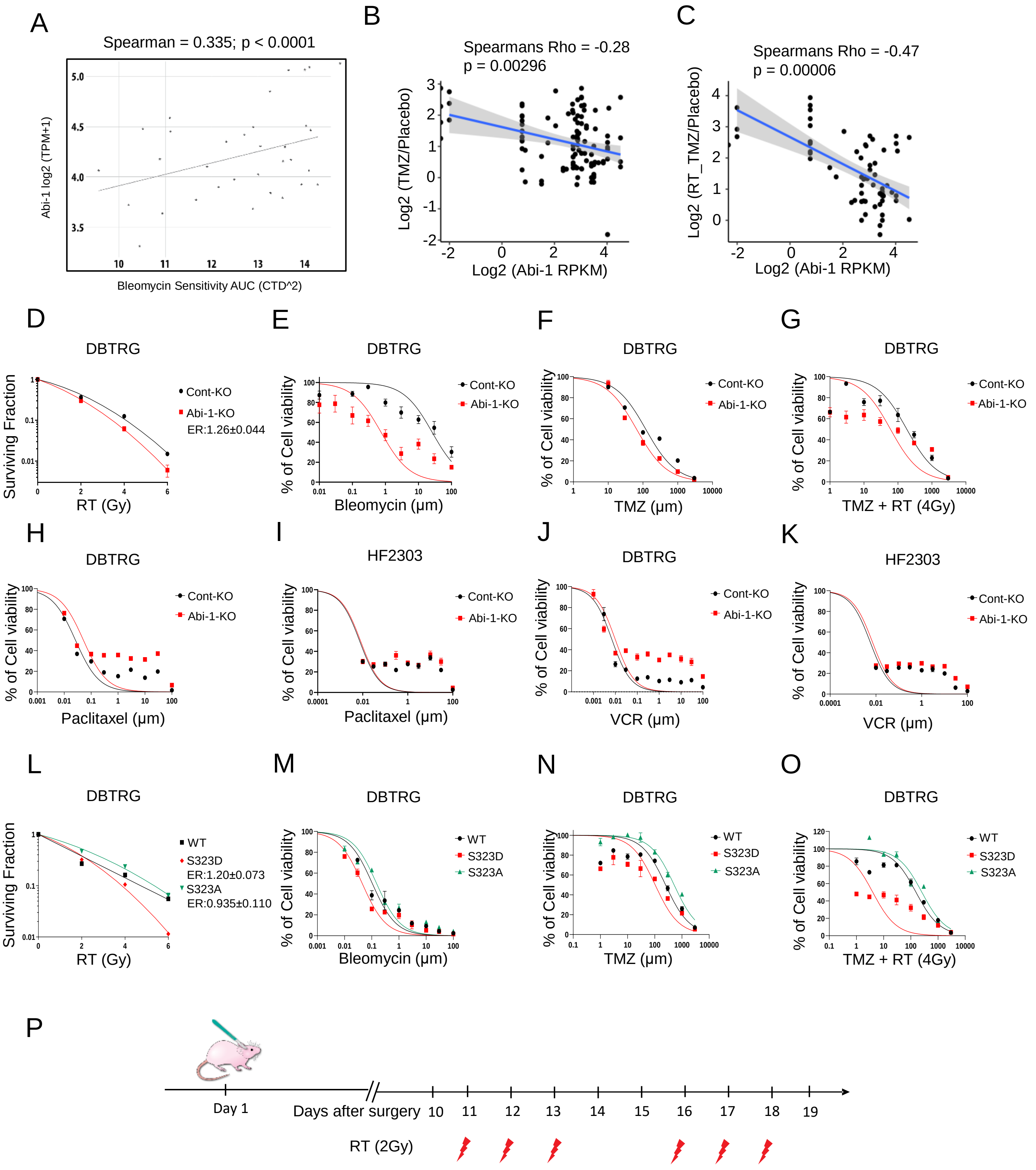

### *Zhou etal.*, Figure S7

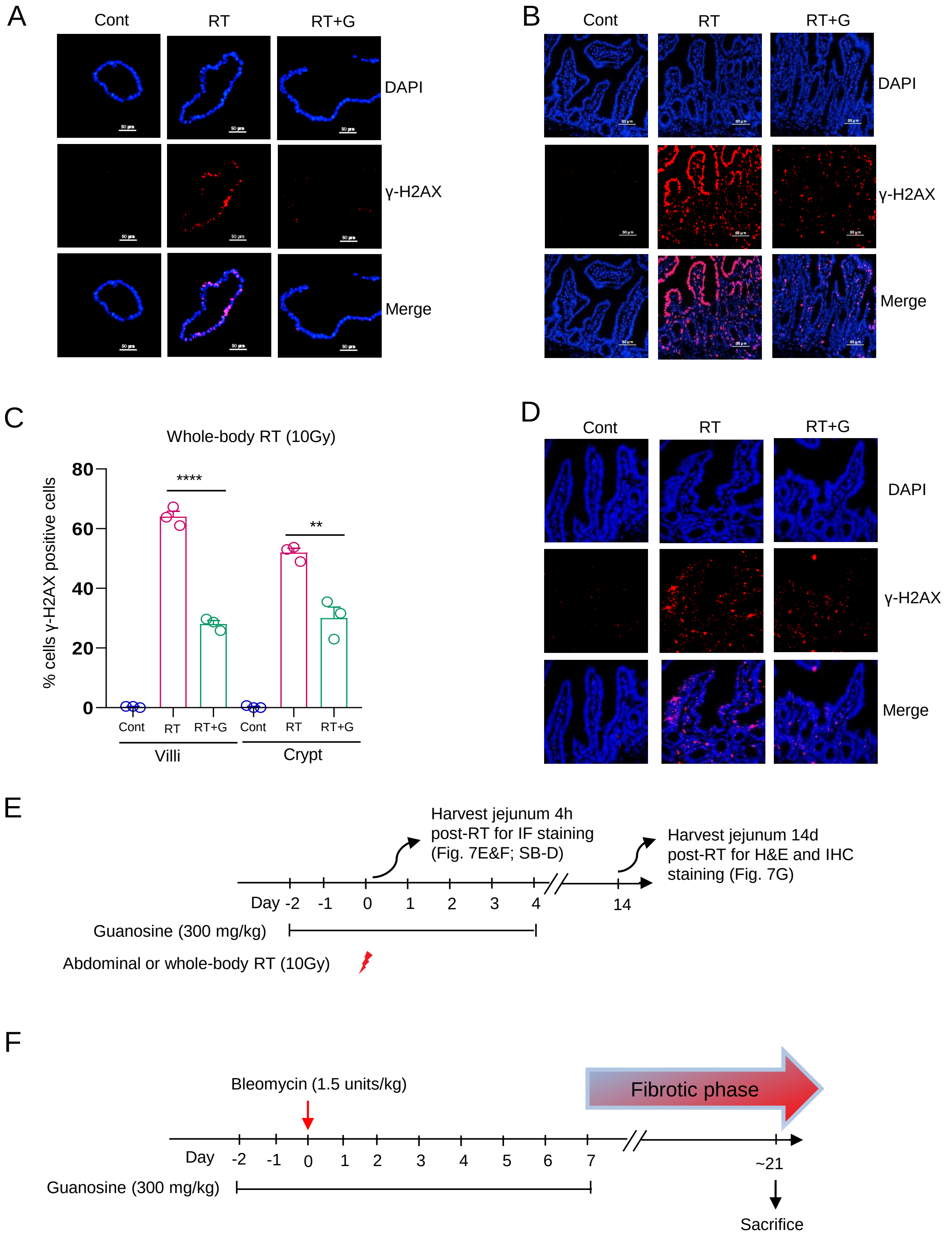
